# Spatially variable competition contributes to mismatched responses of plant fitness and occurrence to environmental gradients

**DOI:** 10.1101/2024.07.25.605199

**Authors:** Kenji T. Hayashi, Nathan J. B. Kraft

## Abstract

1. Species distributions have long been understood to depend on the complex interplay between the abiotic environment and biotic interactions. Empirical work in ecological communities has increasingly revealed how competition can mediate species’ demographic responses to environmental variation, but understanding how the demographic consequences of spatially variable competition manifest in observed distribution patterns remains an important challenge.
2. Here, we describe a conceptual framework for characterizing competition-driven mismatches between responses of fitness and occurrence to environmental gradients. We then explore these mismatches for eight annual plant species in an edaphically variable California grassland landscape.
3. We experimentally quantified how species’ demographic rates (germination rate, fecundity) and fitness respond to spatial variation in the soil environment, either in the presence or absence of naturally occurring neighbors. We also surveyed species’ occurrence across the study landscape. Combining these demographic and occurrence data, we asked whether observed occurrence patterns are congruent with responses of intrinsic fitness (i.e., fitness in the absence of competitors) to soil gradients.
4. We found that competition altered responses of fitness to the primary soil gradient (soil texture) for many (4/8) species. In turn, occurrence patterns were often poorly or even inversely related to responses of intrinsic fitness to this environmental axis. In contrast, we found that competition had relatively little effect on responses of fitness and occurrence to a secondary soil gradient (soil Ca:Mg).
5. *Synthesis*. We demonstrate that spatially variable competition can contribute to mismatched responses of fitness and occurrence to environmental variation. Importantly, these quantitative mismatches depend on the species and environmental gradient in question. Our results caution against assuming that variation in occurrence implies variation in intrinsic fitness (or vice versa) without first disentangling how the abiotic environment and competition impact the demographic processes that underlie species distributions.

## Introduction

Understanding how the abiotic environment and biotic interactions jointly shape species distributions is a long-standing challenge in ecology and biogeography (Dobzhansky 1950; Louthan *et al*. 2015; MacArthur 1972). In particular, establishing whether species’ current distributions reflect their direct demographic responses to the environment, or rather are modified by complex biotic interactions, is a critical step in predicting species’ future distributions (Davis *et al*. 1998; Pearson & Dawson 2003). Although there is widespread theoretical (e.g., Godsoe *et al*. 2017; Price & Kirkpatrick 2009; Svenning *et al*. 2014; Usinowicz & Levine 2018) and empirical (e.g., Armitage & Jones 2020; Legault *et al*. 2020; Usinowicz & Levine 2021; Wisz *et al*. 2013) evidence that biotic interactions such as competition can limit where species are found, quantifying the effects of competition on demography and distributions along real-world environmental gradients is often logistically challenging (but see e.g., Craig *et al*. 2023; Lyu & Alexander 2022; Stanton-Geddes *et al*. 2012). Thus, as environmental change drives shifts in the abundance and distributions of species both within and across ecological communities (e.g., Bowler *et al*. 2017; Chen *et al*. 2011; Feeley *et al*. 2020; Kelly & Goulden 2008; Parmesan & Yohe 2003; Rosenblad *et al*. 2023), competition remains a critical source of uncertainty in forecasting the future of biodiversity (Alexander *et al*. 2016; Gilman *et al*. 2010; HilleRisLambers *et al*. 2013).

The first principles of population dynamics dictate that species persist in environments where they have a positive population growth rate (fitness) when rare (Grainger *et al*. 2019; Hutchinson 1978). Environmentally driven variation in fitness in the absence of biotic interactions (intrinsic fitness) is the foundation for species distributions (Ehrlén & Morris 2015; Holt 2009; Schurr *et al*. 2012), but other processes can create mismatches between intrinsic fitness and observed distributions (Pulliam 2000). For example, competition can prevent species from establishing or persisting in certain environments by reducing fitness in the presence of competitors (realized fitness) to below self-replacement levels (Chesson 2000b; MacArthur & Levins 1967). More subtly, competition can create discrepancies between how intrinsic and realized fitness vary in response to environmental gradients if the identity, density, or per capita effects of competitors depend on the environment (Louthan *et al*. 2015). Notably, recent experimental work has demonstrated not only that the outcomes of competition depend on environmental context (e.g., Cervantes-Loreto *et al*. 2023; Germain *et al*. 2018; Van Dyke *et al*. 2022; Wainwright *et al*. 2019) but also that competition can mediate species’ responses to environmental change (e.g., Esch *et al*. 2018; Liancourt *et al*. 2013). These findings imply that species’ current or future distributions may be incongruent with their intrinsic demographic responses to the environment (Figure 1), which limits our ability to predict species’ responses to environmental change. Progress on this topic requires a better understanding of how spatially variable competition shapes species distributions along real-world environmental gradients.

**Figure 1:**
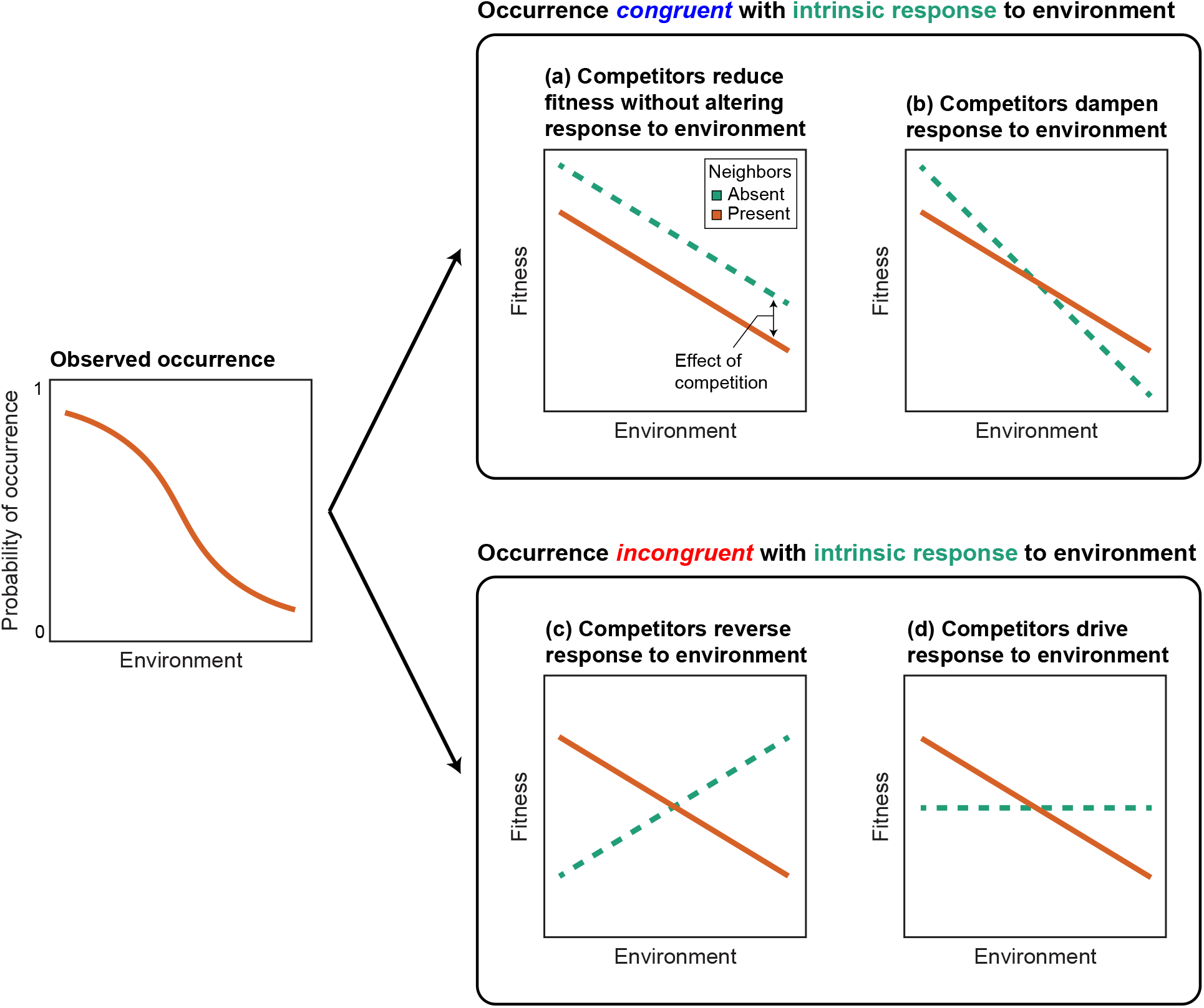
Conceptual framework for identifying competition-driven mismatches between responses of occurrence and fitness to environmental gradients. Observed occurrence patterns (leftmost panel) may be expected to reflect species’ realized demographic responses to the environment (i.e., demographic responses in the presence of competitors; solid orange lines, identical in panels a–d). However, the extent to which observed occurrence patterns track species’ intrinsic demographic responses to the environment (i.e., demographic responses in the absence of biotic interactions; dashed green lines) is often unclear. If competitors (a) uniformly reduce fitness or (b) merely dampen demographic responses, occurrence can be approximately congruent with species’ intrinsic responses to the environment. However, if demographic responses are (c) reversed or (d) even driven by spatially variable competition, occurrence patterns will be incongruent with species’ intrinsic responses to the environment. Note that positive effects of neighbors (i.e., facilitation) are allowed here for graphical simplicity, but are not necessary for these scenarios to play out within a given range of environmental conditions.

A common prediction is that species experience more intense competition in environments where they exhibit higher intrinsic fitness. This idea is foundational for the study of plant strategies (e.g., CSR model, Grime 1977) and species coexistence (Chesson 2000a) in spatially variable environments. Similarly, the stress-gradient hypothesis posits that competition increases in frequency or strength in more benign abiotic environments (Bertness & Callaway 1994; Maestre *et al*. 2009). Such co-variation between intrinsic fitness and competition can emerge if species share functional traits or evolutionary histories that promote similar responses to environmental variation (Adler *et al*. 2013; Mayfield & Levine 2010). While the importance of competition for species distributions has long been hypothesized to depend on environmental context (Dobzhansky 1950; Louthan *et al*. 2015; MacArthur 1972), the consequences of co-variation between intrinsic fitness and competition are rarely studied in the context of species distributions (but see e.g., Armitage & Jones 2020; Usinowicz & Levine 2021). Perhaps counterintuitively, species might be more likely to occur in more competitive environments if competition only weakly dampens demographic responses to the environment, such that occurrence approximately tracks intrinsic fitness (Figure 1b). However, with stronger competition in intrinsically favorable environments, competition can reverse demographic responses to the environment (Figure 1c).

Even in the absence of intrinsic responses, variation in competition can drive realized demographic responses to the environment (Figure 1d). Importantly, these last two scenarios (Figure 1c, d) both create incongruence between trends in observed occurrence and intrinsic fitness along environmental gradients. Beyond the well-established understanding that the absence of a species from an area does not necessarily reflect the suitability of the abiotic environment (Davis *et al*. 1998; Pearson & Dawson 2003), the complexities of these scenarios highlight the need to critically evaluate the practical assumption that species’ distributions approximate their intrinsic demographic responses to environmental variation.

Here, we build up an understanding of how the abiotic environment and competition interact to shape the demography and distributions of species in an experimentally tractable natural landscape. Specifically, we conducted a spatially distributed demographic experiment to evaluate the consequences of spatially variable competition for plant demography and distributions in an edaphically heterogeneous California annual grassland. We experimentally quantified how the demography and fitness of eight annual plant species respond to variation in the abiotic environment, either in the presence or absence of naturally occurring competitors. We asked: (i) What are species’ intrinsic demographic responses to the abiotic environment? (ii) How does competition modify these intrinsic responses? Then, combining these demographic results with occurrence surveys, we asked: (iii) Are observed occurrence patterns congruent with intrinsic or realized demographic responses to the environment? Our results reveal complex and variable relationships between fitness, competition, and occurrence depending on the species and environmental gradient under consideration. Critically, we find that occurrence patterns are often poorly related to—or even in opposition to—species’ intrinsic demographic responses to environmental gradients, highlighting the complex role of spatially variable competition in shaping species distributions.

## Materials and methods

### Study system

We conducted our study at the University of California Natural Reserve System Sedgwick Reserve in Santa Barbara County, California, USA. This region experiences a Mediterranean climate with hot, dry summers and cool, wet winters. We focused on a ∼4-ha area of the reserve characterized by serpentine-derived hummocks interspersed among a grassland matrix (Figure 2a, Figure S1.1). The hummocks host many native annual forbs, whereas the matrix is dominated by invasive annual grasses (e.g., *Avena* spp., *Bromus* spp.) (Gram *et al*. 2004). The abiotic and biotic heterogeneity of this landscape, coupled with the dominance of experimentally tractable annual plants, makes this system well-suited for addressing our research questions.

**Figure 2:**
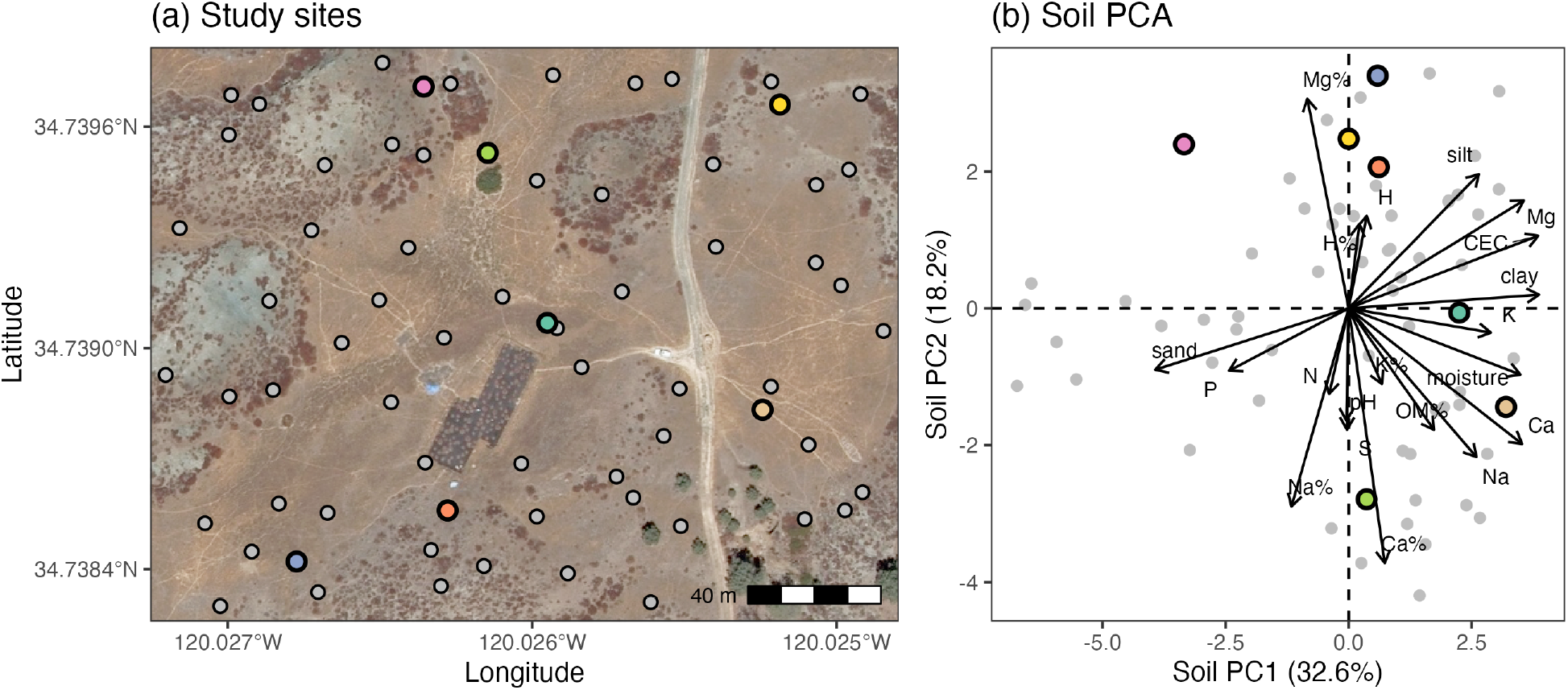
Locations and soil environments of study sites. Colored points represent experimental sites. Gray points represent survey sites. (a) Map of study sites. High resolution orthoimagery courtesy of the U.S. Geological Survey. (b) Principal component analysis (PCA) of soil variables at study sites. Arrows represent the loadings of individual soil variables with respect to PC1 and PC2. The percentage of variance explained by each axis is given in parentheses.

### Demographic experiment

To quantify the demographic responses of annual plant species to spatial variation in the environment and competition, we grew eight native annual plant species (Table 1) at seven sites distributed across the study area (Figure 2a). These experimental sites represent a subset of sites previously selected to capture environmental variation across this area (Kandlikar *et al*. 2022). Each site (∼4 m × ∼4 m) was comprised of 20 0.5 m × 0.5 m plots, evenly divided among two neighborhood treatments in which the resident community was either removed or left intact prior to planting. Each plot contained eight 15 cm × 15 cm subplots, each of which was seeded at its center with a fixed number of seeds (24 to 40 depending on the species) of one of the eight focal species in November 2019. Seeds were collected and aggregated from across the study area in 2016–2018.

**Table 1:** Plant species used in this study.

| Code | Species | Family |
| --- | --- | --- |
| ACWR | *Acmispon wrangelianus* | Fabaceae |
| CHGL | *Chaenactis glabriuscula* | Asteraceae |
| FEMI | *Festuca microstachys* | Poaceae |
| HECO | *Hemizonia congesta* | Asteraceae |
| LACA | *Lasthenia californica* | Asteraceae |
| PLER | *Plantago erecta* | Plantaginaceae |
| SACO | *Salvia columbariae* | Lamiaceae |
| URLI | *Uropappus lindleyi* | Asteraceae |

In January–February 2020, we recorded germination rate (*g*) in each subplot by counting the number of germinants found in the seeded locations, then thinned these germinants down to a single focal individual. We marked focal individuals with toothpicks and tracked them for the duration of the growing season. Due to below-average rainfall in January–February (County of Santa Barbara 2024), we added ∼0.6 in of water to all plots following germination, equivalent to a typical storm at this time of year. In April–July 2020, as focal individuals started to show signs of senescence, we estimated the lifetime fecundity (*F*) of each individual. See Appendix S2 for detailed methods for estimating fecundity. In total, we recorded germination rate in 1,078 subplots and estimated fecundity for 708 focal individuals. We maintained the neighborhood removal treatment by weeding out any background germinants throughout the experiment.

### Occurrence surveys

To characterize our focal species’ distributions across the study landscape, we revisited all experimental sites (colored points in Figure 2) in April–May 2021 and surveyed the occurrence of each focal species in each uncleared experimental plot. We also surveyed occurrence at 61 additional sites distributed across the study area (gray points in Figure 2). We selected these survey sites by overlaying on the study area a 7 × 9 grid of 25 m × 25 m cells and randomly generating a point within each grid cell using QGIS (QGIS Development Team 2020). Each survey site was comprised of a single 0.5 m × 0.5 m plot, equivalent to one experimental plot.

### Soil sampling

To characterize the soil environment at our study sites, we collected soil samples at all sites (seven experimental sites + 61 survey sites) alongside occurrence surveys. We collected soil from ∼5–10 cm below the soil surface at three locations within each site (one near the center and two along the edges of the site). We aggregated soil samples for each site and submitted them to A&L Western Laboratories in Modesto, California, USA for chemical and physical analysis. We similarly collected soil samples following a rainfall event and estimated soil moisture as gravimetric water content. See Table S1.1 for descriptions of all soil variables used in this study.

### Statistical analyses

To describe the soil environment at our study sites, we performed a principal component analysis (PCA) of the 20 soil variables in Table S1.1 (Figure 2b, Figure S1.2, Figure S1.3). Based on results of parallel analysis (Dinno 2024; Horn 1965), we retained the first four axes (adjusted eigenvalues > 1) for all subsequent analyses. These axes correspond to 77% of cumulative variance explained (PC1 = 32.6%, PC2 = 18.2%, PC3 = 16.9%, PC4 = 9.3%).

We used Bayesian hierarchical models to estimate the demographic responses of our focal species to the soil environment in the presence or absence of naturally occurring neighbors. For germination rate, we modeled the number of germinants as following a beta-binomial distribution with an expected probability of germination, *μ*, and precision, *ϕ*. We specified a linear model for *μ* with a logit link function as:

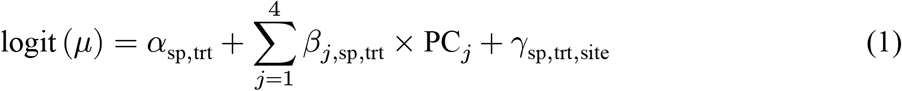

where *α* is an intercept and *β*_*j*_ is a slope for soil principal component axis *j* = 1, …, 4 for each species (sp = 1, …, 8) and neighborhood treatment (trt = 1 is with neighbors present; trt = 2 is with neighbors absent). For clarity, we hereafter refer to the slope coefficients in the presence or absence of neighbors as *β*_present_ and *β*_absent_, respectively. We also apply this convention to other quantities estimated in the presence or absence of neighbors. *γ* is a random intercept for each species and treatment at each site = 1, …, 7. *γ* was hierarchically modeled as:

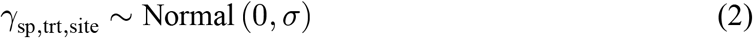

where *σ* is the standard deviation that describes variation in *γ*. Next, we modeled fecundity as following a zero-inflated negative binomial distribution, which combines (i) a negative binomial distribution with an expected fecundity, *μ*, and shape, *ϕ*, with (ii) an expected probability of zero-inflation, *θ*. We specified linear models for *μ* and *θ* in the same form as the right-hand side of Equation 1, using a log link function for *μ* and a logit link function for *θ*. For both models, we allowed *ϕ* to differ by species. See Appendix S3 for a full account of these models.

To estimate the response of fitness to the soil environment and competition, we implemented a joint model of the germination and fecundity models described above. We then used the joint posterior distribution for the germination and fecundity sub-models to compute fitness as the finite rate of increase (*r*) in the functional form of a model of seed banking annual plants (Chesson 1990; MacDonald & Watkinson 1981):

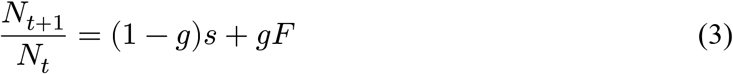

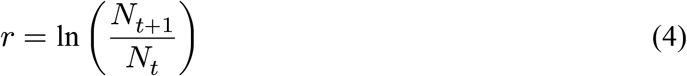

where *N*_*t*_ is the number of seeds in year *t* and *s* is the annual rate of seed survival in the seed bank. We used species-specific estimates of *s* compiled from previous work in this study area (Godoy *et al*. 2014, 2017; Levine & HilleRisLambers 2009), assuming *s* to be constant across sites. We employed this joint modeling approach, rather than directly modeling fitness, because excess zeros in our fecundity data contributed to biomdalities in fitness estimates computed from our experimental data. We used this joint fitness model for all demographic analyses. In the context of our demographic experiment, we refer to fitness estimated in the presence or absence of neighbors as realized fitness (*r*_present_) and intrinsic fitness (*r*_absent_), respectively.

We used a Bayesian generalized linear model to quantify the occurrence patterns of our focal species along soil gradients. To align the scope of this analysis with our demographic experiment, we focused on occurrence data for sites within the range of soil PC1–PC4 captured by experimental sites, corresponding to 92 plots at 31 sites. We modeled occurrence as following a beta-binomial distribution with an expected probability of occurrence, *μ*, and precision, *ϕ*. We specified a linear model for *μ* following Equation 1, but excluding the random intercept term (*γ*) and without indexing other terms by neighborhood treatment (trt) as these components were not applicable here. We allowed *ϕ* to differ by species. See Appendix S4 for a full account of this model. Then, to evaluate the congruence between occurrence and fitness along soil gradients, we computed the posterior probability that the slopes (*β*_*j*_) for occurrence and fitness in response to each of soil PC1–PC4 have the same sign. That is, for each posterior sample, we asked whether the slopes for occurrence and fitness (in the presence or absence of neighbors) along a given soil axis were both positive or both negative, then calculated the proportion of posterior samples that satisfied this criterion. Note that the expected response of fitness to soil PC1–PC4 was nonlinear, so we computed an axis-wide slope 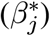 by taking the difference in expected fitness at maximum (max(PC_*j*_)) and minimum values of the axis (min(PC_*j*_)) and dividing this difference by ΔPC_*j*_ = | max(PC_*j*_) *−* min(PC_*j*_)|.

We implemented Bayesian statistical models using Stan (Stan Development Team 2024) via cmdstanr (Gabry *et al*. 2024). For all models, we used weakly informative priors that were intended to keep parameters within plausible ranges given nonlinear link functions (Wesner & Pomeranz 2021) and prior work in our study system (Godoy *et al*. 2014; Kandlikar *et al*. 2022; Kraft *et al*. 2015; Van Dyke *et al*. 2022). We confirmed model convergence by inspecting diagnostic quantities 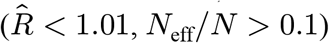 and trace plots. We report posterior support for statistical effects as the probability of direction (i.e., the probability of a positive or negative effect, [0.5, 1]) where applicable. We also report credible intervals as 95% highest-density continuous intervals (HDCIs) unless stated otherwise. See Appendices S3 and S4 for details of our statistical modeling approach. We conducted analyses using R version 4.4.1 (R Core Team 2024).

## Results

### Soil environment

Soil PC1 was characterized by a gradient in soil texture, with clay content loading positively and sand content loading negatively along the axis (Figure 2b). Cation exchange capacity (CEC), cation content, and soil moisture positively co-varied with clay content along soil PC1. As such, this axis also captures important differences in soil fertility. Soil PC2 was characterized by a gradient in cation composition, with Mg saturation loading positively and Ca saturation loading negatively along the axis (Figure 2b). All sites exhibited an excess of Mg relative to Ca (Ca:Mg < 1) characteristic of serpentine soils (Brady *et al*. 2005; Fernandez-Going *et al*. 2012). Taken together, our study sites capture continuous variation in the canonical physical and chemical properties of serpentine soils (Gram *et al*. 2004; Walker 1954). Therefore, we focus on results for soil PC1 and PC2 below. Soil PC3 and PC4 appeared to reflect variation in pH and organic matter content, respectively (Figure S1.2).

### Demographic responses to environment and competition

Our focal species tended to exhibit higher intrinsic demographic performance (i.e., demographic performance in the absence of neighbors) in more fine-textured soils with higher Mg saturation. In the absence of neighbors, germination rate increased with soil PC1 or PC2 for 3/8 species (Pr(*β*_absent_ > 0) ≥ 0.975; Figure S3.2, Figure S3.3, Table S3.2). Responses of fecundity to soil PC1 and PC2 were less clear (two species with Pr(direction) ≥ 0.95; Figure S3.4, Figure S3.5, Table S3.5). Combining these demographic rates, fitness increased with soil PC1 or PC2 for 5/8 species 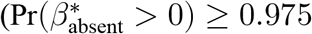 Figure 3, Figure S3.6, Table S3.8).

**Figure 3:**
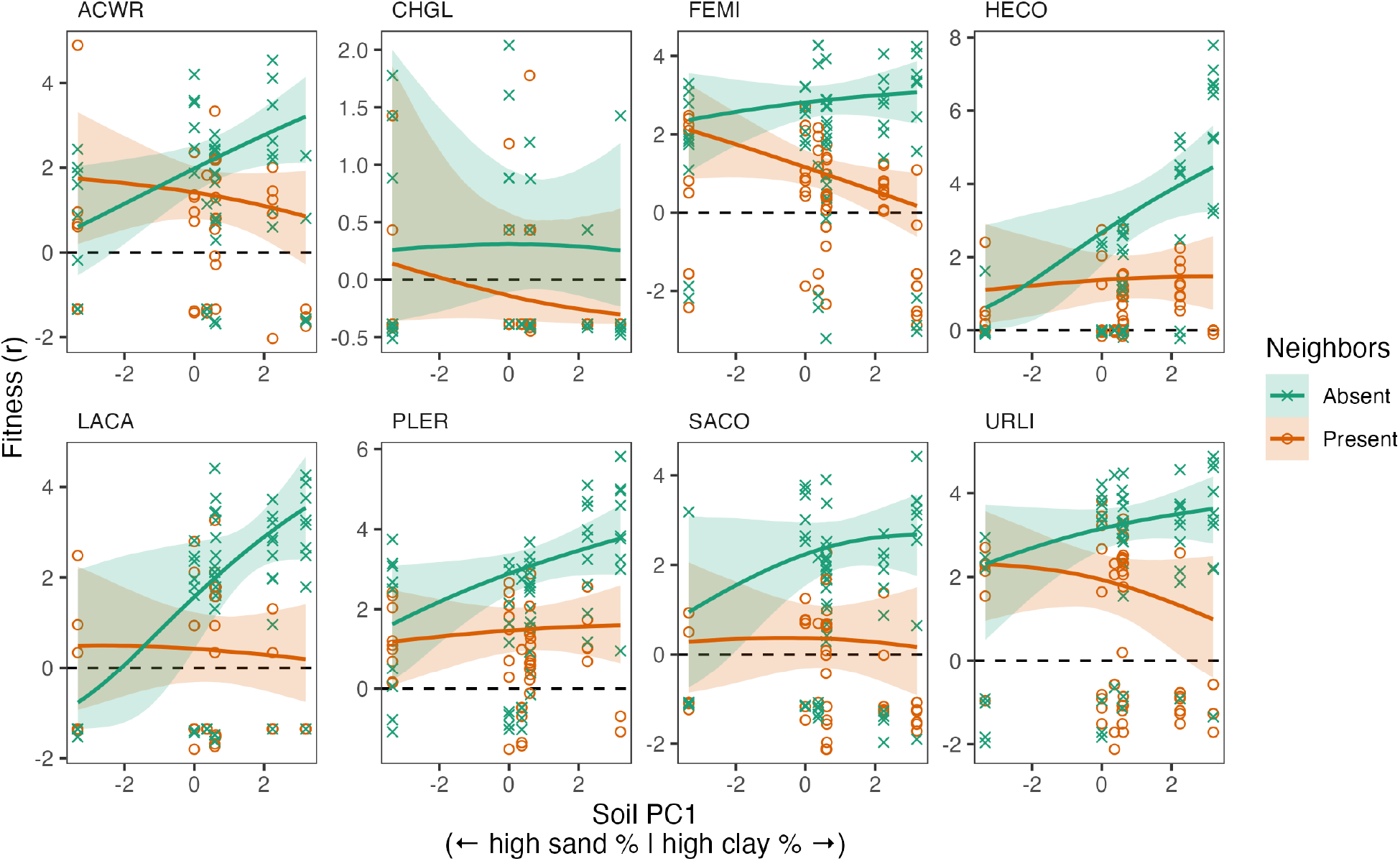
Response of fitness (*r*) to soil PC1 in the presence or absence of neighbors. Posterior expectations are shown for each species with soil PC2–PC4 held at average conditions across experimental sites. Solid lines represent medians. Shaded areas represent 95% quantile intervals.

Neighbors had variable effects on fitness across environments. On average, neighbors decreased the fitness of all but one of our focal species (Pr(Δ*r* < 0) ≥ 0.975, where Δ*r* = *r*_present_ *− r*_absent_; Figure S3.1, Table S3.1). Additionally, we found evidence that neighbors alter the response of fitness to soil PC1 (i.e., there is an interaction effect between neighborhood treatment and soil PC1, 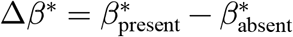). Specifically, neighbors reduced the slope for fitness in response to soil PC1 for 4/8 species: ACWR and LACA (Pr(Δ*β*^*^ < 0) ≥ 0.975), as well as FEMI and HECO with marginally less support (Pr(Δ*β*^*^ < 0) > 0.972; 95% HDCIs do not overlap 0) (Figure 3, Table S3.10). For ACWR, HECO, and LACA, neighbors flattened the response of fitness to soil PC1, wherein positive intrinsic responses to this axis were largely negated by the presence of neighbors (Table S3.8; Table S3.9). Notably, for FEMI, neighbors appeared to largely drive the realized response of fitness to soil PC1 (Table S3.8; Table S3.9), consistent with Figure 1d. In contrast, we did not find clear evidence that neighbors alter the response of fitness to soil PC2 (Pr(direction) < 0.884 for all species; Figure S3.6, Table S3.10). Overall, we found contrasting roles of competition in mediating demographic responses to the soil environment, with competition altering responses of fitness to a soil texture gradient (PC1) while having relatively minimal effect on responses of fitness to a soil nutrient (Ca:Mg) gradient (PC2).

Results for germination rate and fecundity are shown in Appendix S3.

### Congruence between occurrence and fitness

In accordance with our demographic results, the degree of congruence between trends in occurrence and fitness along soil gradients was highly variable depending on the species and soil axis under consideration. Occurrence was negatively associated with soil PC1 for 5/8 species (Pr(*β* < 0) ≥ 0.975; Figure 4, Table S4.1) and positively associated with soil PC2 for one species (Pr(*β* > 0) ≥ 0.975; Figure S4.1, Table S4.1). Strikingly, for all but two species, the responses of occurrence and intrinsic fitness to soil PC1 were more likely than not to be in opposite directions (Figure 5; mean probability of congruence = 0.27, SE = 0.074). For instance, LACA is expected to exhibit a positive intrinsic response to soil PC1 (Figure 3), yet its probability of occurrence is expected to decrease along this axis (Figure 4), resulting in a probability of congruence of 0.021. In contrast, the responses of occurrence and intrinsic fitness to soil PC2 tended more toward being congruent with one another (Figure S4.2; mean probability of congruence = 0.60, SE = 0.094). Accounting for the effects of neighbors on fitness tended to improve congruence between occurrence and fitness, especially with respect to soil PC1 (Figure 5; mean probability of congruence = 0.64, SE = 0.075). These results provide empirical support for spatially variable competition as an important but complex driver of mismatches between occurrence and fitness along environmental gradients.

**Figure 4:**
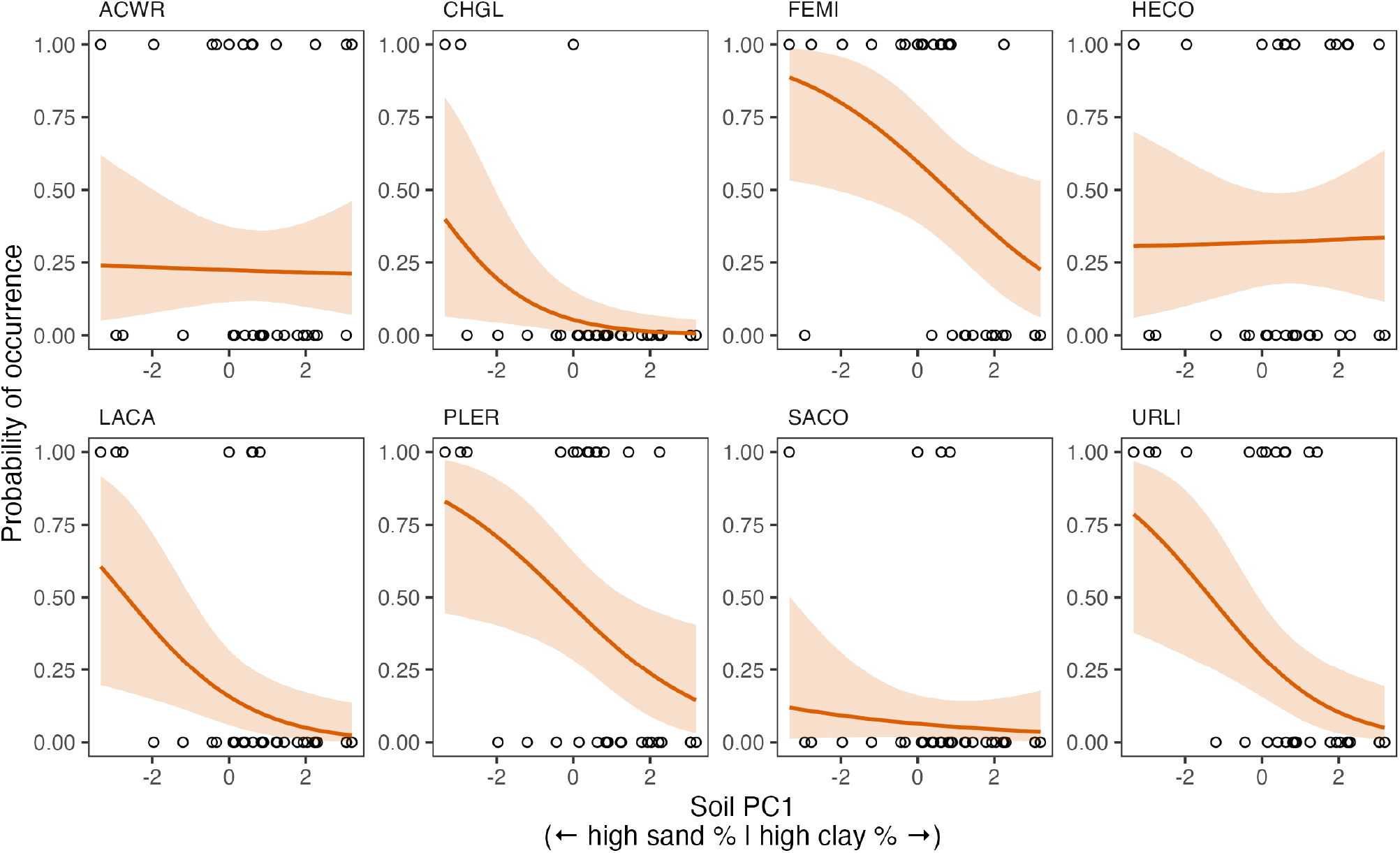
Observed patterns of occurrence along soil PC1. Posterior expectations are shown for each species with soil PC2–PC4 held at average conditions across sites. Solid lines represent medians. Shaded areas represent 95% quantile intervals.

**Figure 5:**
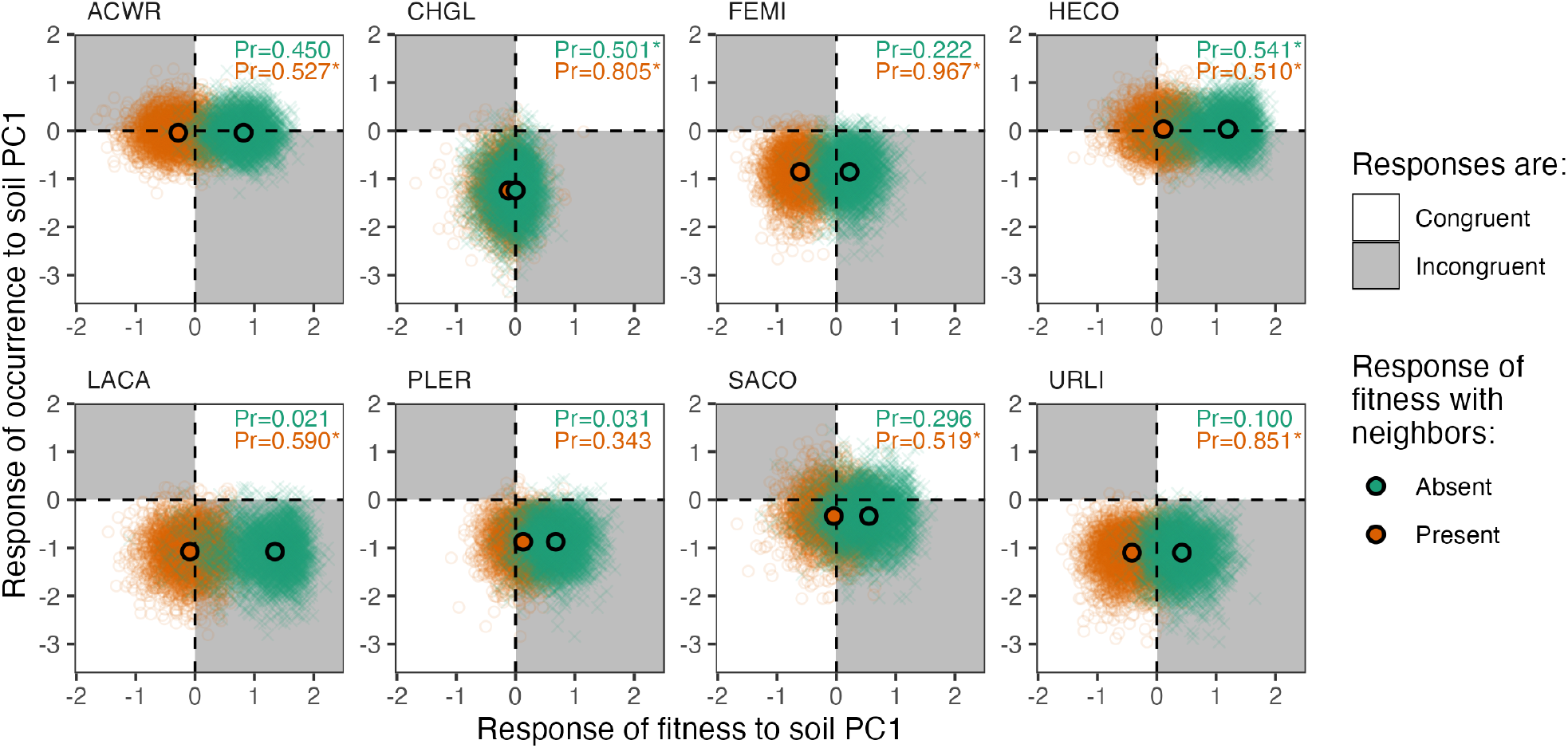
Congruence between responses of occurrence and fitness to soil PC1. For each species, the slope of occurrence in response to soil PC1 (*β*_*j*=1_ for the occurrence model) is plotted against the axis-wide slope of fitness in response to soil PC1 (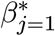 for the fitness model), the latter of which is computed in the presence (orange) or absence (green) of neighbors. Point clouds are posterior samples (thinned to 4,000 samples for visualization) and center points are medians. White regions denote congruent responses (i.e., slopes are both positive or both negative) and gray regions denote incongruent responses (i.e., one slope is positive while the other slope is negative). The total probability of congruence (i.e., the proportion of posterior samples that fall in the white regions) is shown in the top-right corner of each panel. Asterisks indicate that responses are more likely to be congruent that not (i.e., Pr(congruence) > 0.5); note that these are not results of statistical significance tests.

## Discussion

Competitive interactions have long been understood to contribute to species distributions (Wisz *et al*. 2013), from the zoning of barnacles in the rocky intertidal (Connell 1961) to the migration of plant species in response to climate change (Alexander *et al*. 2015). While empirical work in ecological communities has increasingly revealed how competition can mediate species’ demographic responses to environmental variation (e.g., Cervantes-Loreto *et al*. 2023; Esch *et al*. 2018; Germain *et al*. 2018; Liancourt *et al*. 2013; Van Dyke *et al*. 2022; Wainwright *et al*. 2019), understanding how the demographic consequences of competition manifest in observed distribution patterns remains challenging. Disentangling the contributions of intrinsic fitness (i.e., fitness in the absence of biotic interactions) and competition to species distributions in natural landscapes is an important step toward more accurately predicting species distributions (Davis *et al*. 1998), especially as environmental change threatens to modify biotic interactions in ecological communities (Alexander *et al*. 2016; Gilman *et al*. 2010; HilleRisLambers *et al*. 2013). Here, we combined a spatially distributed demographic experiment with occurrence surveys to evaluate the consequences of competition for plant species’ demography and distributions in a California annual grassland. We found evidence that competition can alter the response of fitness to environmental gradients, effectively decoupling realized fitness (i.e., fitness in the presence of neighbors) from intrinsic fitness. Furthermore, we found that competition often contributes to incongruence between occurrence and fitness, such that observed occurrence patterns can be poorly or even inversely related to species’ intrinsic responses to environmental variation. Notably, competition had contrasting effects on fitness and occurrence depending on the species and environmental gradient under consideration. Our findings illustrate the importance of accounting for spatially variable competition in the study of species distributions and caution against the practical but simplifying assumption that species’ distributions track their intrinsic demographic responses to environmental variation.

Competition is most often described as limiting species distributions by excluding species from intrinsically suitable environments (e.g., Bertness & Ellison 1987; Bullock *et al*. 2000; Connell 1961). Accordingly, much research has focused on the role of competition in setting discrete distributional limits at the edges of species’ ranges (e.g., Anderegg & HilleRisLambers 2019; Ettinger & HilleRisLambers 2013; Louthan *et al*. 2015; Lyu & Alexander 2022; Price & Kirkpatrick 2009; Stanton-Geddes *et al*. 2012). However, much of the ecological world is not well described by discrete transitions. Species often exhibit continuous variation in demographic performance in response to continuous variation in the abiotic and biotic environment. Therefore, understanding how competition drives quantitative variation in occurrence (or abundance) is critical to disentangling the processes shaping species’ current and future distributions. In particular, an important way that competition can influence distribution patterns is by altering the response of fitness to the abiotic environment (Figure 1).

In our experiment, we found that competitors altered several (4/8) species’ fitness responses to a soil texture gradient (PC1; Figure 3). In turn, we found that observed occurrence patterns often failed to track responses of intrinsic fitness to this axis (Figure 5). In fact, most (6/8) species were more likely than not to exhibit an increased probability of occurrence in less intrinsically favorable conditions along soil PC1. Such mismatched responses of occurrence and intrinsic fitness to soil PC1 were especially prominent for LACA and PLER (Pr(congruence) < 0.032), both of which are forb species often associated with the transition zone between serpentine hummocks and grassland matrix (Gram *et al*. 2004). In contrast, competitors had relatively little effect on species’ fitness responses to a soil nutrient (Ca:Mg) gradient (PC2; Figure S3.6), and observed occurrence patterns were more likely to be congruent with intrinsic responses of fitness to this axis (Figure S4.2). Our results expand on previous findings that competition can alter species’ demographic responses to environmental variation (e.g., Esch *et al*. 2018; Liancourt *et al*. 2013) by providing empirical evidence that such altered responses can quantitatively impact species distributions. Importantly, we show that competition can even contribute to trends in occurrence along environmental gradients that are opposite to what would be expected from demographic responses in the absence of competitors.

Our results have particularly important implications for predicting or interpreting species distributions based on observed distribution patterns. Expanding on earlier work (Davis *et al*. 1998; Pearson & Dawson 2003), we find that competition not only contributes to incongruence between occurrence and fitness, but also has contrasting consequences depending on the species and environmental gradient in question. This complexity cautions against extrapolating from observed distribution patterns without careful consideration of how particular species experience competition along specific environmental gradients. In the absence of detailed demographic data, widely utilized species distribution models (SDMs) can account for competition by including proxies of competitive effects (e.g., population density) as predictors (Elith & Leathwick 2009; Wisz *et al*. 2013). The variable effects of competition on demography and distributions observed in this study suggest that SDMs should ideally allow the effects of competition proxies to vary by species and to interact with environmental variables. These implications also extend to demographically or physiologically informed models of species’ niches and distributions (e.g., Benito Garzón *et al*. 2019; Ehrlén & Morris 2015; Kearney & Porter 2009; Merow *et al*. 2014; Schurr *et al*. 2012), wherein inadequately accounting for variable effects of competition might yield misleading predictions in terms of how species’ distributions are expected to track their intrinsic responses to environmental gradients.

Our demographic results were partially consistent with long-standing predictions of co-variation between intrinsic fitness and competition across spatial environmental gradients (Figure 1b, c) (Bertness & Callaway 1994; Chesson 2000a; Grime 1977). In particular, 3/8 species (ACWR, HECO, LACA) displayed higher intrinsic fitness in finer-textured soils (high PC1), where they were also more limited by competition. Consequently, responses of fitness to soil PC1 for these species were largely flattened by the presence of competitors (Figure 3). Although evaluating the mechanisms underlying variation in competition was beyond the scope of this study, this pattern could be driven in part by an increased dominance of invasive grasses such as *Avena fatua, Bromus diandrus*, and *Bromus hordeaceus* in sites with finer-textured soils (Figure S1.4). These invasive species have been shown to suppress native plants in California serpentine grasslands through a variety of mechanisms, including resource competition, recruitment limitation, and habitat modification (e.g., Chen *et al*. 2018; Hamilton *et al*. 1999; LaForgia 2021; Seabloom *et al*. 2003a, b). As such, the competitive effects measured in our study may be viewed as integrating over multiple types of negative effects imposed by neighbors. While this approach has the benefit of capturing realistic, total effects of interactions, dissecting different sources of variation in competitive effects is an important next step toward a more mechanistic understanding of species distributions (see also Louthan *et al*. 2015).

Considering that our demographic experiment captured only a single generation of population dynamics in a system with substantial interannual rainfall variability (Levine & Rees 2004), albeit in a water year with about average total rainfall (2019–2020 = 21.57 in + 0.6 in added manually, mean = 21.44 in, County of Santa Barbara 2024), it is notable that we were able to detect imprints of competition on observed occurrence patterns across our study landscape. However, similar to previous work that found occurrence and demographic parameters to be weakly associated (Thuiller *et al*. 2014), observed occurrence patterns often could not be clearly explained by experimentally estimated responses of fitness to the soil environment and competition (Figure 5; Figure S4.2). One possibility is that interannual variation in the competitive environment introduces complexities in the population dynamics that underlie occurrence (e.g., Hallett *et al*. 2019; Hobbs *et al*. 2007; Pitt & Heady 1978). Additional processes such as dispersal limitation, source-sink dynamics, demographic stochasticity, and other biotic interactions may also contribute to observed occurrence patterns (Pulliam 2000). For example, Craig *et al*. (2023) found that source-sink dynamics play an integral role in decoupling annual plant occupancy from fitness in a Northern California grassland at both the species and community level. We note that, despite an abundance of sites where one or more of our focal species were absent (Figure 4, Figure S4.1), we only found up to moderate (∼80%) support for environmental filtering (i.e., absence from sites due to *r*_absent_ < 0, sensu Kraft *et al*. 2015) and competitive exclusion (i.e., absence from sites due to *r*_present_ < 0 when *r*_absent_ > 0) along soil PC1 and PC2 in our experiment (Figure S3.7, Figure S3.8). This observation aligns with previous findings that the (re-)establishment of native plants in California grasslands is often seed-limited (Germain *et al*. 2017; Seabloom *et al*. 2003a, b), but more work is needed to explore this possibility in our study landscape.

## Conclusions

The complex interplay between demography and competition across abiotic environments is a critical source of uncertainty in explaining or predicting species distributions. In particular, understanding how the demographic consequences of competition translate to species distributions along real-world environmental gradients remains an important challenge. Here, we outlined a conceptual framework for identifying quantitative mismatches between fitness and occurrence along environmental gradients. We then provided experimental evidence for mismatched responses of plant fitness and occurrence to edaphic gradients in a California annual grassland. Importantly, we showed that competition can contribute to observed trends in fitness and occurrence along environmental gradients that are decoupled—or even reversed—from responses of intrinsic fitness (i.e., fitness in the absence of neighbors) to environmental variation. However, the consequences of competition for fitness and occurrence were highly dependent on the species and environmental gradient under consideration. Our results caution against assuming that observed patterns of fitness or occurrence track species’ intrinsic responses to environmental variation without careful consideration of spatially variable competition. Targeted demographic studies such as ours can complement statistical species distribution models by revealing when and how observed distribution patterns reflect intrinsic fitness vs. competitive interactions.

## Acknowledgements

We acknowledge the Chumash peoples as the traditional land caretakers of the area where we conducted our field study. We acknowledge the Gabrielino/Tongva peoples as the traditional land caretakers of Tovaangar (Los Angeles basin, So. Channel Islands), where UCLA is located. We thank T. Foster, G. Kandlikar, E. Purington, and M. Vaz for assistance with the field experiment. We thank Sedgwick Reserve staff, including L. Johnsen and K. McCurdy, for support at the reserve. We are grateful to A. Cisneros Carey, L. Glevanik, H. Oyler, M. Sagarin, K. Schneider, M. Tingley, and J. Yanowitz for feedback on the manuscript. The manuscript also benefited from discussion with A. Angert and C. Kremer. This work was performed at the University of California Natural Reserve System Sedgwick Reserve (https://doi.org/10.21973/N3C08R) and supported by a Mildred E. Mathias Graduate Student Research Grant from the University of California Natural Reserve System. This work was supported by National Science Foundation grants DEB 1644641 and DEB 2022810.

## Author contributions

KTH and NJBK conceptualized the study and developed the methodology. KTH led investigation, data curation, and formal analysis with input from NJBK. KTH led writing of the original draft of the manuscript with substantial input from NJBK. KTH and NJBK contributed to review and editing of the manuscript.

## Data availability statement

Original data and code for analyses are archived on Zenodo at https://doi.org/10.5281/zenodo.12978683 (Hayashi & Kraft 2024).

## Conflict of interest statement

The authors have no conflicts of interest to declare.

## Notes

### Competing Interest Statement

The authors have declared no competing interest.

### Summary of Updates

Revised statistical analyses. Revised title, main text, figures, and supplementary materials accordingly. Most notably, Figure 1, Figure 5, and associated text were refined.

