## Supplementary materials for "Spatially variable competition contributes to mismatched responses of plant fitness and occurrence to environmental gradients"

Kenji T. Hayashi Nathan J. B. Kraft

### Appendix S1: Study landscape and soil environment

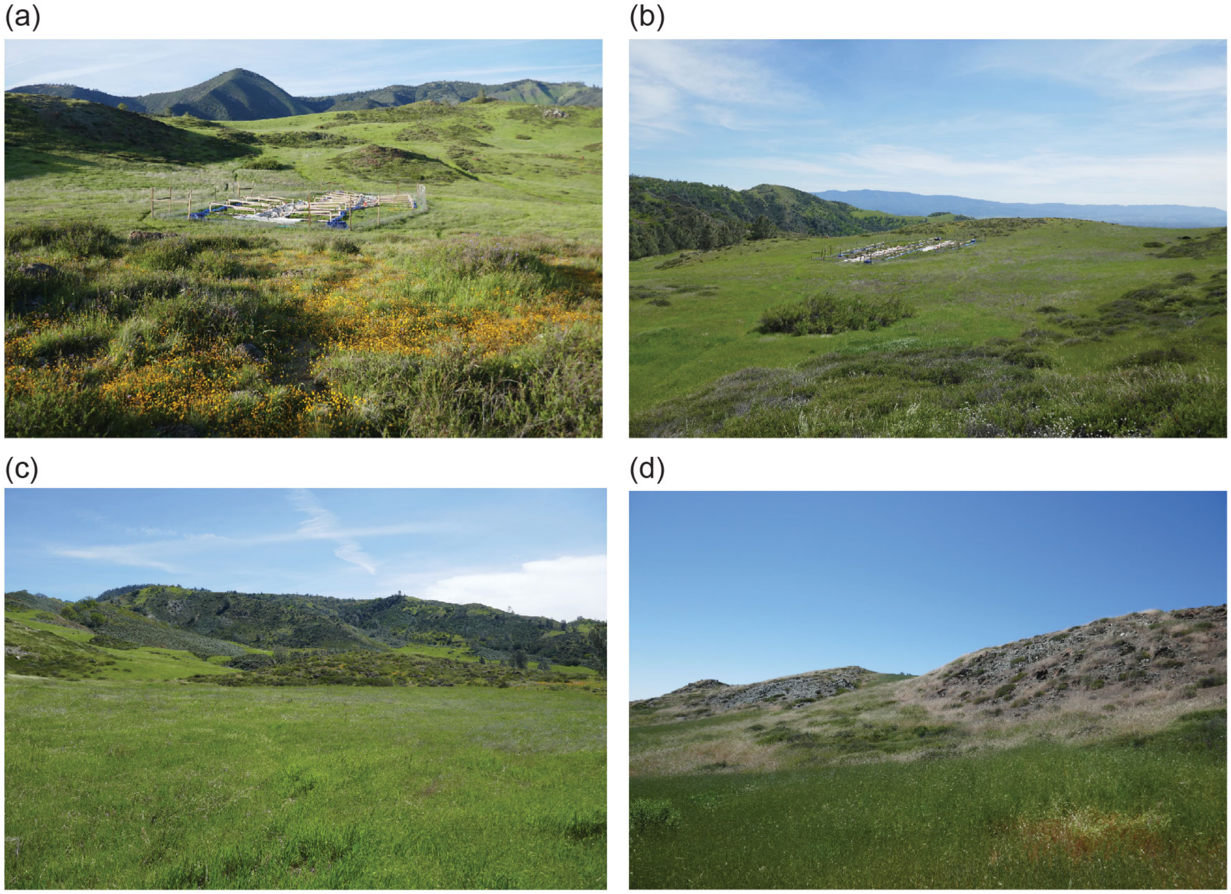

Figure S1.1: Photographs of the study landscape taken facing (a) north, (b) south, (c) east, and (d) west in April 2019. Note that the enclosure visible in (a, b) is not part of this study. Photo credit: Kenji Hayashi.

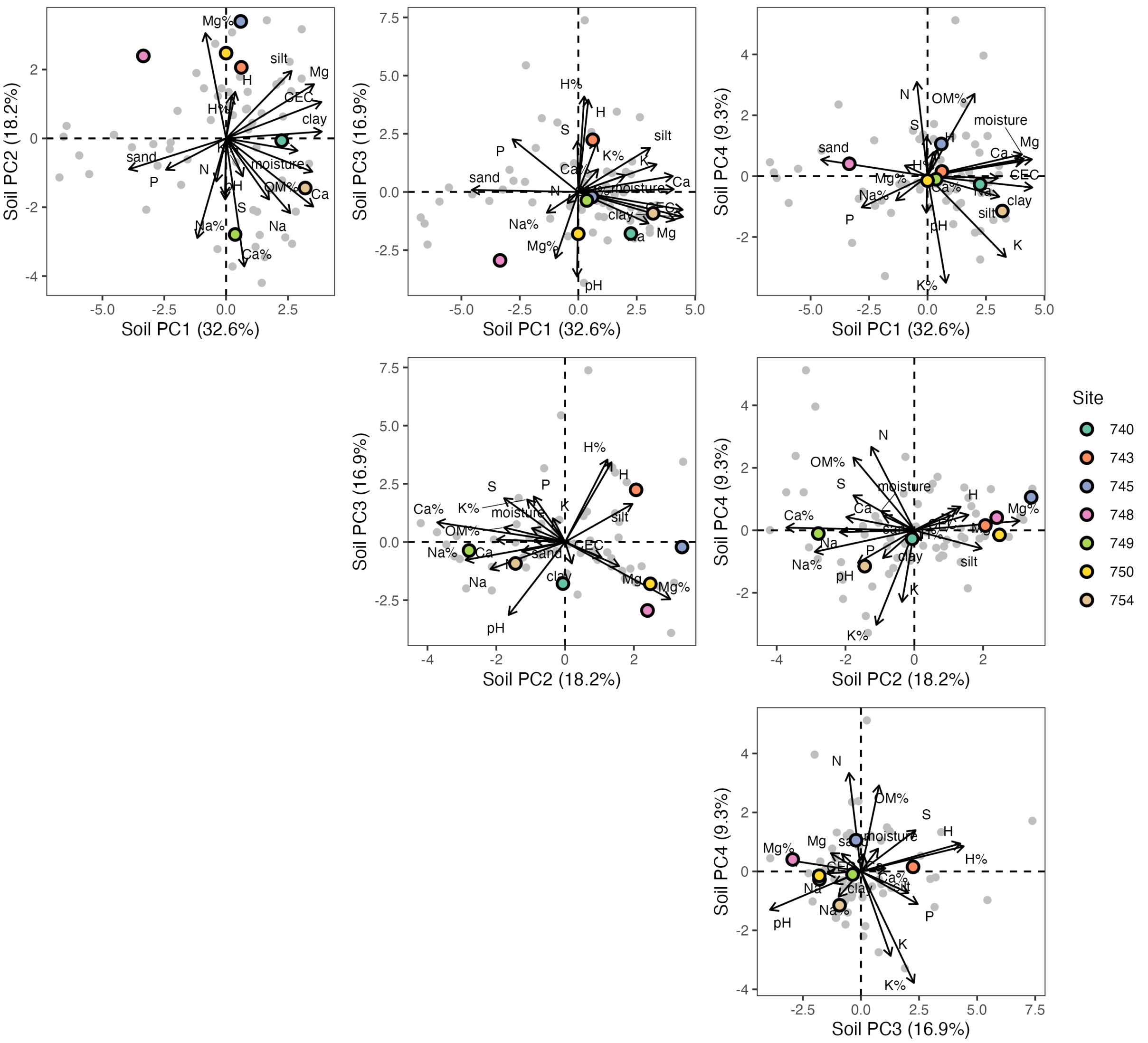

Figure S1.2: Principal component analysis (PCA) of soil variables at study sites. The first four axes, corresponding to 77% of cumulative variance explained, are shown here. Colored points represent experimental sites. Gray points represent survey sites. Arrows represent the loadings of individual soil variables with respect to each axis. The percentage of variance explained by each axis is given in parentheses.

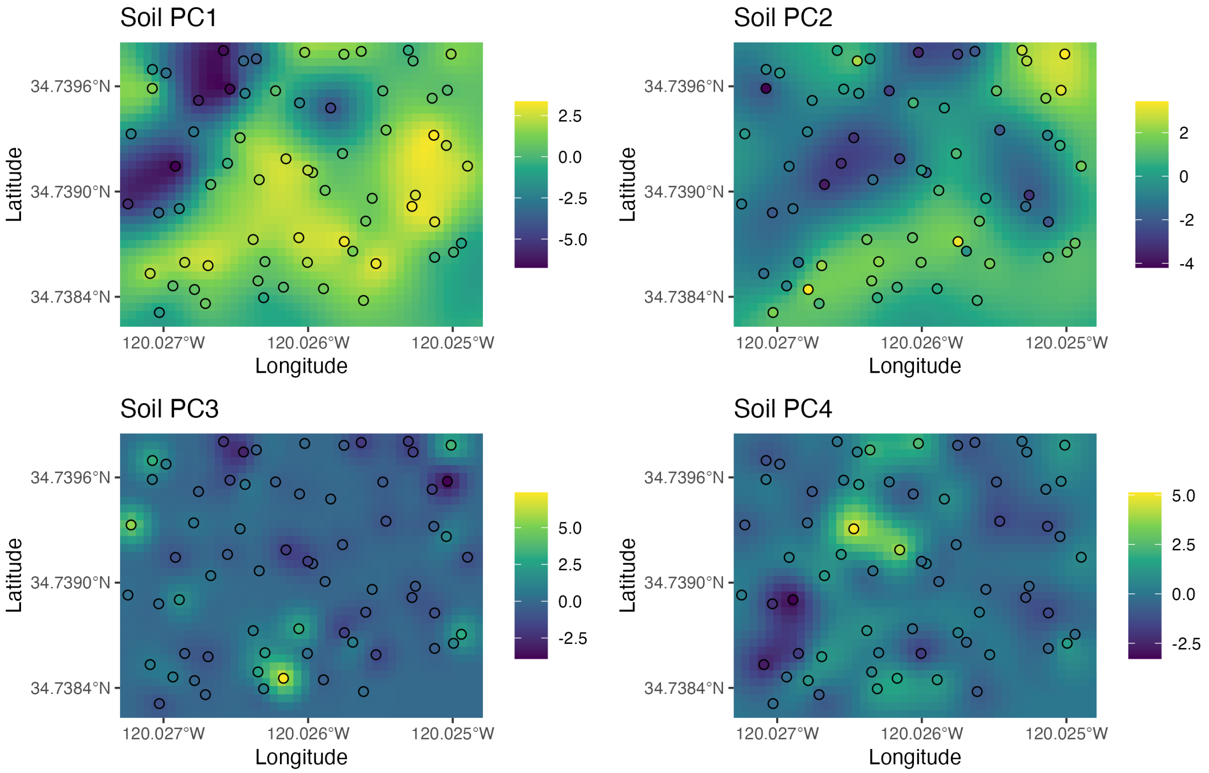

Figure S1.3: Interpolated maps of soil principal component axes for the study landscape. Maps were generated by performing ordinary kriging with a spherical variogram model using `gstat` (Gräler *et al.* 2016; Pebesma 2004). Points represent the sites at which soil samples were collected and are colored according to their principal component scores.

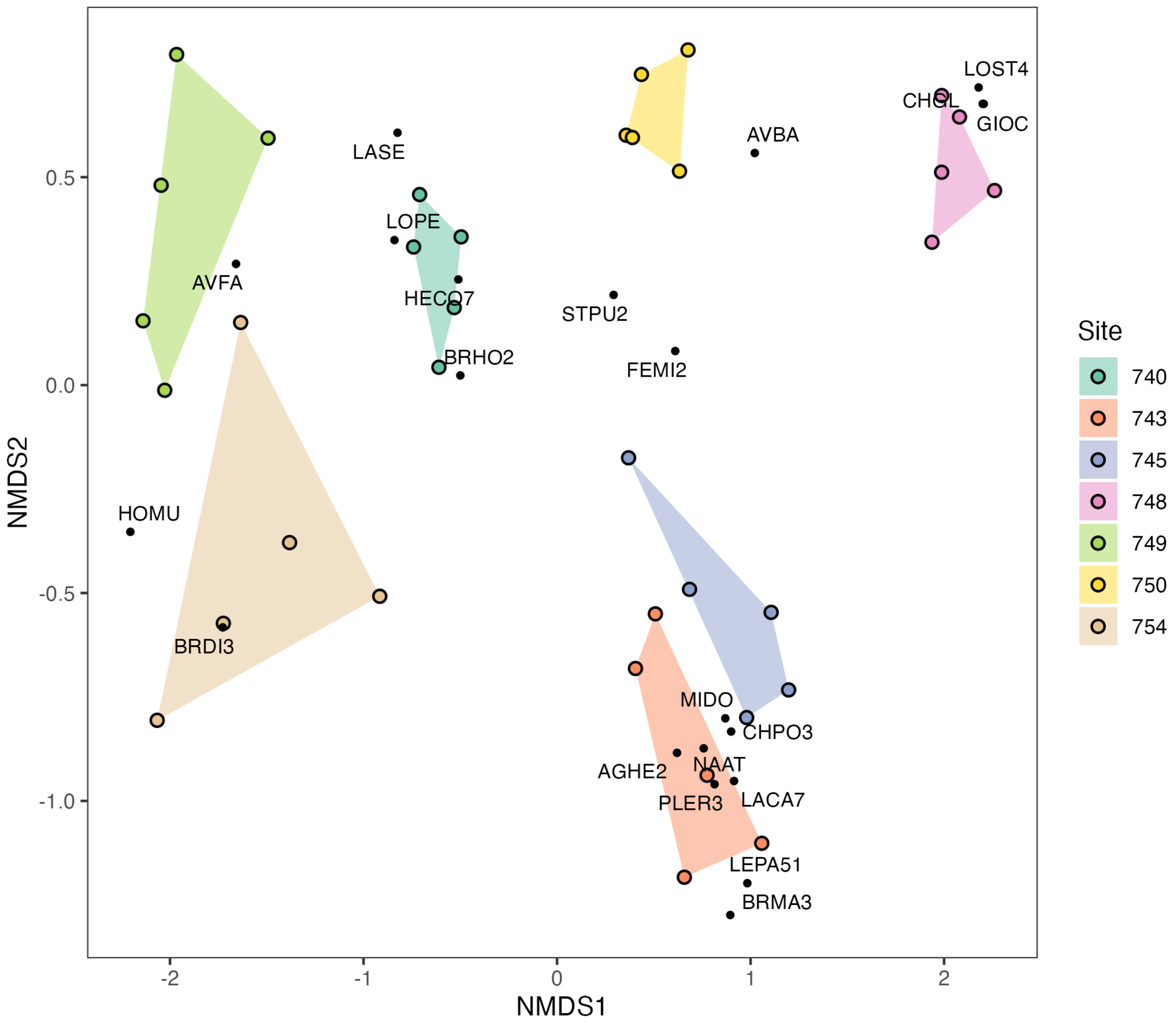

Figure S1.4: Nonmetric multidimensional scaling (NMDS) for plant species cover at experimental sites. NMDS was performed using `vegan` (Oksanen *et al.* 2024) with Bray-Curtis dissimilarity and $k$ (number of dimensions) = 2. Cover data were collected at our experimental sites in 2017 (data from Kandlikar 2021; Kandlikar *et al.* 2022). Colored points correspond to replicate plots at each site. Black points correspond to the (expanded) weighted average scores for species with over 5% cover in any plot. Species are labeled with their symbols in the U.S. Department of Agriculture PLANTS Database.

Table S1.1: Soil variables used in principal component analysis.

| Variable | Description | Units |
|---|---|---|
| OM% | Organic matter content | % |
| P | Phosphorus content | ppm |
| K | Potassium content | ppm |
| Mg | Magnesium content | ppm |
| Ca | Calcium content | ppm |
| Na | Sodium content | ppm |
| H | Hydrogen content | meq/100g |
| pH | pH | none |
| CEC | Cation exchange capacity | meq/100g |
| K% | Potassium saturation | % |
| Mg% | Magnesium saturation | % |
| Ca% | Calcium saturation | % |
| Na% | Sodium saturation | % |
| H% | Hydrogen saturation | % |
| N | Nitrate-nitrogen ($NO_3$-N) content | ppm |
| S | Sulfate-sulfer ($SO_4$-S) content | ppm |
| sand | Sand content | % |
| silt | Silt content | % |
| clay | Clay content | % |
| moisture | Gravimetric water content | % |

### Appendix S2: Methods for estimating fecundity

We estimated the lifetime fecundity of focal individuals in our demographic experiment by counting the number of seeds produced by each individual. However, direct seed counts in the field were impractical for some species. For ACWR, CHGL, HECO, LACA, and URLI, we first counted the number of reproductive structures on each focal individual. For ACWR, we counted the number of seed pods. For CHGL, HECO, LACA, and URLI, we counted the number of inflorescences. Next, we collected and dissected ~40 of these reproductive structures for each species and calculated the mean number of seeds per reproductive structure (Figure S2.1). Finally, we estimated fecundity for each focal individual by multiplying the number of reproductive structures by the mean number of seeds per reproductive structure.

For SACO, we measured the radius of all seed heads (approximated as circles) on each focal individual. We collected and dissected ~40 of these seed heads and counted the number of seeds per seed head. We then fit a negative binomial generalized linear model with the number of seeds per seed head as the response and seed head radius as the predictor (Figure S2.2). We implemented this model using the `glm.nb` function in `MASS` (Venables & Ripley 2002) with the formula `seeds ~ radius`, where `seeds` is the number of seeds per seed head and `radius` is seed head radius. We used a negative binomial likelihood in anticipation of overdispersion that could arise from collecting seed heads from across sites. We used this model to predict the expected number of seeds for each seed head, then estimated fecundity for each focal individual as the sum of the expected number of seeds for all of its seed heads.

We also considered a model in which seed head area was used as the predictor of the number of seeds per seed head for SACO (formula: `seeds ~ area`, where `area` is seed head area). In both models, the predictor was centered and scaled to unit variance. We used Akaike Information Criterion (AIC) to compare alternative models and found that the radius model was estimated to have better predictive performance (Table S2.1). Therefore, we used the radius model to estimate fecundity for SACO as described above.

For PLER, we counted the number of flowers (or fruits), where each flower (or fruit) was observed to produce two seeds. For FEMI, we directly counted the number of seeds. All fecundity estimates were rounded down to integer values (where applicable). All counts of seeds and reproductive structures included those that were still emerging or developing and traces of those that were already lost at the time of fecundity estimation. Fecundity for marked focal individuals that were confirmed or inferred to have experienced mortality prior to seed production were recorded as 0. For subplots in which no focal individuals were marked (e.g., due to germination failure) or focal individuals were lost (e.g., due to disturbances such as gopher damage), fecundity was recorded as missing and excluded from analyses. Following estimation of fecundity, we removed focal individuals from plots to minimize seed set from these individuals.

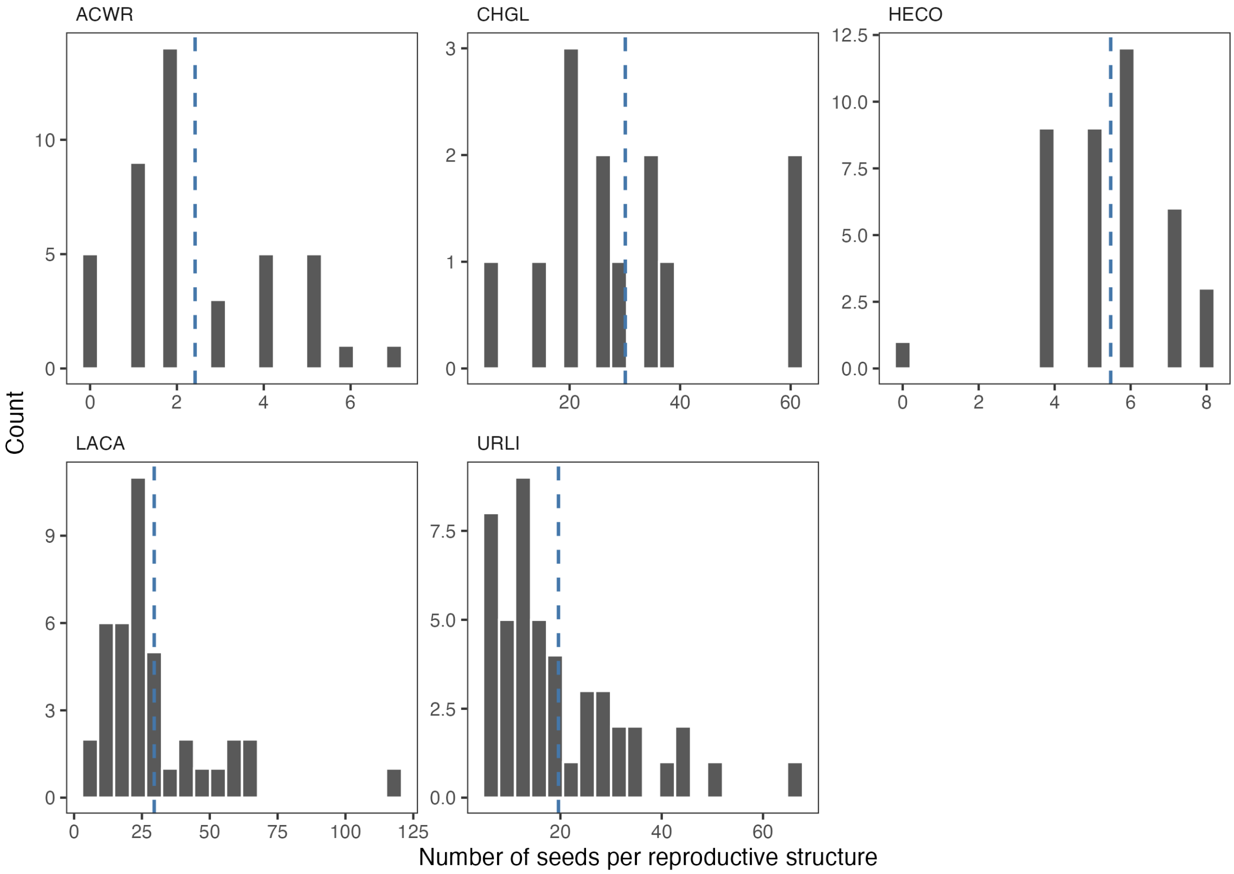

Figure S2.1: Histograms of the number of seeds per reproductive structure for ACWR, CHGL, HECO, LACA, and URLI. Vertical dashed lines represent the mean value for each species.

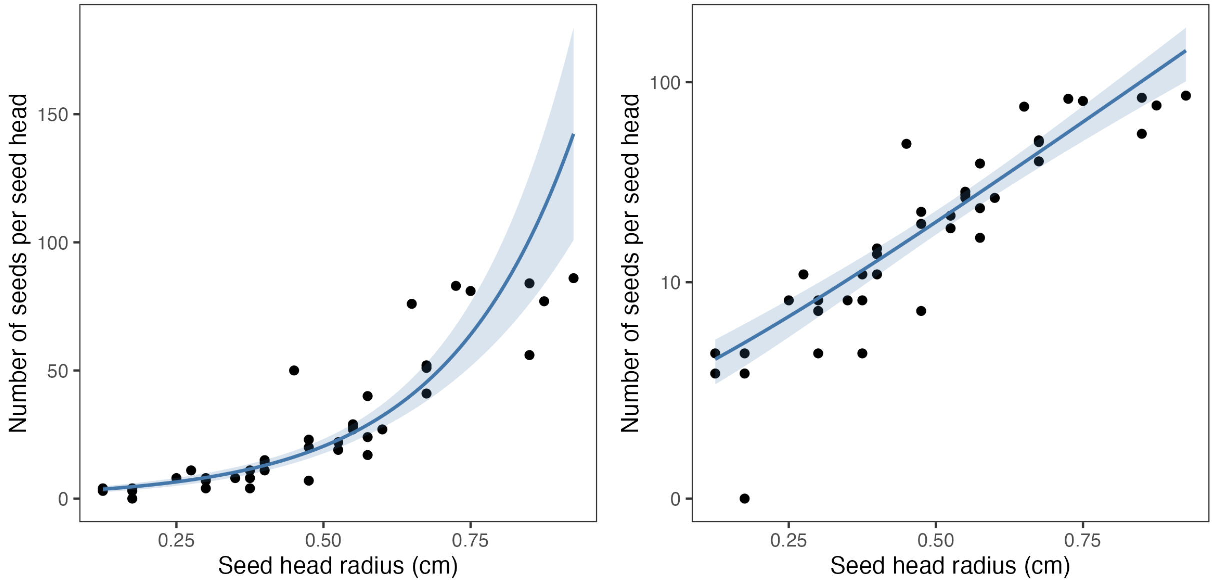

Figure S2.2: Relationship between seed head radius and the number of seeds per seed head for SACO. The relationship is shown on both identity (left) and log (right) scales. Solid lines represent expected values. Shaded areas represent $\pm$ 1.96 SE around the expected values.

Table S2.1: Comparison of models for the number of seeds per seed head for SACO. We used Akaike Information Criterion (AIC) to compare models with alternative predictors. The model selected for use in analyses is indicated with bold text.

| Model | Predictor | AIC | ΔAIC |
|---|---|---|---|
| **(2)** | **Radius** | **285.89** | **0.00** |
| (1) | Area | 302.60 | 16.71 |

### Appendix S3: Analysis of demographic experiment

We used Bayesian hierarchical models to estimate the demographic responses of our focal species to the soil environment and competition. We started by fitting separate models for germination rate ($g$) and fecundity ($F$), which we then implemented jointly as a single model. We used this joint model, which we refer to as the *fitness model*, for all demographic analyses. We implemented all Bayesian models using Stan (Stan Development Team 2024) via `cmdstanr` (Gabry *et al.* 2024).

#### Germination model

We modeled the number of germinated seeds ($n_i$) for observation $i$ = 1, …, 1,078 as following a beta-binomial distribution:

$$n_i \sim \text{Beta-Binomial}\left(N_i, \mu_i, \phi_i\right) \tag{1}$$

where $N$ is the number of planted seeds, $\mu$ is the mean probability parameter, and $\phi$ is the precision parameter. This parameterization of the beta-binomial distribution follows that employed by `brms` (Bürkner 2017). Using a logit link function, we defined $\mu$ as:

$$\text{logit}\left(\mu_i\right) = \alpha_{\text{sp}_i,\text{trt}_i} + \sum_{j=1}^{4} \beta_{j,\text{sp}_i,\text{trt}_i} \times \text{PC}_{j,i} + \gamma_{\text{sp}_i,\text{trt}_i,\text{site}_i} \tag{2}$$

where $\alpha$ is an intercept and $\beta_j$ is a slope for soil principal component axis $j$ = 1, …, 4. $\alpha$ and $\beta_j$ vary by species (sp = 1, …, 8) and neighborhood treatment (trt = 1, 2). That is, this linear model describes the response of expected germination rate ($\mu$) to soil PC1–PC4 for each species in the presence (trt = 1) or absence (trt = 2) of neighbors. $\gamma$ is a group-level (or 'random') intercept that represents a site-specific deviation for each combination of species and treatment from the corresponding trend given by $\alpha$ and $\beta_j$. $\gamma$ reflects our experimental design, wherein repeated measurements of $\{n, N\}$ were taken for each species and treatment at each site = 1, …, 7. $\gamma$ was hierarchically modeled as:

$$\gamma_{\text{sp,trt,site}} \sim \text{Normal}\left(0, \sigma\right) \tag{3}$$

where $\sigma$ is the standard deviation parameter that describes variation in $\gamma$. Finally, we allowed $\phi$ to vary by species as:

$$\phi_i = \alpha_{\phi,\text{sp}_i} \tag{4}$$

where $\alpha_\phi$ is a species-specific intercept for $\phi$.

We specified prior distributions for parameters as:

$$\alpha \sim \text{Normal}(0, 2) \tag{5}$$

$$\beta_j \sim \text{Normal}(0, 1) \tag{6}$$

$$\sigma \sim \text{Exponential}(1) \tag{7}$$

$$\alpha_\phi \sim \text{Exponential}(1) \tag{8}$$

Equation 5, defined on the logit scale, translates to the probability scale as a roughly flat prior that assigns similar density to most values in $(0, 1)$. Equation 6 was employed as a weakly regularizing prior that avoids assigning much density to strong relationships. Using (weakly) informative priors can be important for generalized linear (mixed) models with nonlinear (e.g., logit, log) link functions, where non-informative priors can imply nonsensical expectations (Wesner & Pomeranz 2021). Note that, for model fitting, raw values of soil PC1–PC4 (let these be $x_j$) were centered and scaled with respect to the mean ($\bar{x}_j$) and standard deviation ($\sigma_{x_j}$) of each axis across experimental sites: $\text{PC}_j = (x_j - \bar{x}_j)/\sigma_{x_j}$. For example, Equation 6 implies that $\beta_j \approx -2$ and $\beta_j \approx 2$ correspond to around the 2.5% and 97.5% quantiles of this prior distribution, respectively. These slopes allow for nearly the entire possible range of expected germination rate ($\mu$) to be traversed in response to four standard deviations of change along just a single axis (e.g., $\text{logit}^{-1}(-4) \approx 0.02$, $\text{logit}^{-1}(4) \approx 0.98$). Equation 7 is a positive-constrained prior that implies variation in $\gamma$ that is of similar magnitude as the expectations defined by Equations 5 and 6. Equation 8 was chosen as a vague, positive-constrained prior for the species-specific precision parameter ($\phi$). R code for conducting prior predictive simulations (Gabry *et al.* 2019; Wesner & Pomeranz 2021) is included as part of the archived data and code for this manuscript (Hayashi & Kraft 2024).

We also considered a model in which a binomial likelihood function was employed. We used approximate leave-one-out cross-validation (LOO-CV) (Vehtari *et al.* 2017), implemented with `loo` (Vehtari *et al.* 2024a), to compare models based on their estimated out-of-sample predictive performance. We applied a moment matching correction as needed (specifically, if the Pareto $k$ value for Pareto-smoothed importance sampling > 0.7) to improve the reliability of LOO-CV results (Paananen *et al.* 2021; Vehtari *et al.* 2024b). We considered models to differ in predictive performance if $|\Delta\text{ELPD}_{\text{LOO}}|/\text{SE}_{\Delta\text{ELPD}_{\text{LOO}}} > 2$, where $\Delta\text{ELPD}_{\text{LOO}}$ is the pairwise difference in the LOO estimate of expected log pointwise predictive density and $\text{SE}_{\Delta\text{ELPD}_{\text{LOO}}}$ is the standard error of this difference (see also Sivula *et al.* 2023). Here, we found that the beta-binomial model (described above) was estimated to have better predictive performance (Table S3.11).

We fit all models using Stan's main Markov chain Monte Carlo sampling algorithm. For each model, we ran four Markov chains in parallel, each with 10,000 iterations (5,000 warmup, 5,000 sampling), yielding 20,000 posterior samples per model. This resulted in an effective sample size (ESS) of at least 10,000 for most parameters, which is the minimum ESS recommended by

Kruschke (2015) for computing credible intervals at the 95% level. We used default options for the initial values and control parameters of the sampler. We assessed model convergence by first checking that the sampler produced no critical runtime warnings (e.g., divergent transitions after warmup). We then checked whether diagnostic quantities satisfied recommended criteria. In particular, we confirmed that the $\widehat{R}$ convergence diagnostic < 1.01 and the ratio of effective sample size to total sample size ($N_{\text{eff}}/N$) > 0.1 for all parameters (Stan Development Team 2022; Vehtari *et al.* 2021). Additionally, we inspected trace plots to confirm that Markov chains were well-mixed and stationary after warmup. We also conducted graphical posterior predictive checks (Gabry *et al.* 2019) to assess model fit.

**Fecundity model**

We modeled fecundity ($F_i$) for observation $i$ = 1, …, 708 as following a zero-inflated negative binomial distribution:

If $F_i = 0$:

$$\text{Pr}\left(F_i \mid \theta_i, \mu_i, \phi_i\right) = \theta_i + (1 - \theta_i) \times \text{NegBinomial}\left(0 \mid \mu_i, \phi_i\right) \tag{9}$$

If $F_i > 0$:

$$\text{Pr}\left(F_i \mid \theta_i, \mu_i, \phi_i\right) = (1 - \theta_i) \times \text{NegBinomial}\left(F_i \mid \mu_i, \phi_i\right) \tag{10}$$

where $\theta$ is the probability of observing $F = 0$ from a zero-inflation process, and conversely, $(1 - \theta)$ is the probability of observing $F \geq 0$ from a negative binomial process. $\mu$ is the mean parameter and $\phi$ is the shape parameter for the negative binomial distribution. Using a log link function, we defined $\mu$ as:

$$\log\left(\mu_i\right) = \alpha_{\text{sp}_i,\text{trt}_i} + \sum_{j=1}^{4} \beta_{j,\text{sp}_i,\text{trt}_i} \times \text{PC}_{j,i} + \gamma_{\text{sp}_i,\text{trt}_i,\text{site}_i} \tag{11}$$

$$\gamma_{\text{sp,trt,site}} \sim \text{Normal}\left(0, \sigma\right) \tag{12}$$

Similarly, using a logit link function, we defined $\theta$ as:

$$\text{logit}\left(\theta_i\right) = \alpha_{\theta,\text{sp}_i,\text{trt}_i} + \sum_{j=1}^{4} \beta_{\theta,j,\text{sp}_i,\text{trt}_i} \times \text{PC}_{j,i} + \gamma_{\theta,\text{sp}_i,\text{trt}_i,\text{site}_i} \tag{13}$$

$$\gamma_{\theta,\text{sp,trt,site}} \sim \text{Normal}\left(0, \sigma_\theta\right) \tag{14}$$

The right-hand sides of Equations 11 and 13 are specified identically to the right-hand side of Equation 2, where the subscript $\theta$ denotes parameters for the zero-inflation component. That is, expected fecundity from the negative binomial process ($\mu$) and the expected probability of zero-inflation ($\theta$) are both allowed to vary for each species and treatment in response to soil PC1–PC4. We allowed $\phi$ to vary by species as:

$$\phi_i = \alpha_{\phi,\text{sp}_i} \tag{15}$$

where $\alpha_\phi$ is a species-specific intercept for $\phi$.

We specified prior distributions for parameters as:

$$\alpha \sim \text{Normal}(4, 2) \tag{16}$$

$$\alpha_\theta \sim \text{Normal}(0, 2) \tag{17}$$

$$\beta_j, \beta_{\theta,j} \sim \text{Normal}(0, 1) \tag{18}$$

$$\sigma, \sigma_\theta \sim \text{Exponential}(1) \tag{19}$$

$$\alpha_\phi \sim \text{Exponential}(1) \tag{20}$$

These priors were specified following a similar rationale as described for the germination model with the exception of Equation 16, which is an informative prior based on previous fecundity estimates for our focal species in our study system (Godoy *et al.* 2014; Kandlikar *et al.* 2022; Kraft *et al.* 2015; Van Dyke *et al.* 2022). Equation 16 implies for example that, under average environmental conditions across experimental sites (i.e., $\text{PC}_j = 0$), an expected fecundity ($\mu$) of $\exp(8) \approx 3{,}000$ corresponds to around the 97.5% quantile of this prior distribution. From this intercept, $\mu$ can be further increased (or decreased) at each site according to $\beta_j$ and $\gamma$. We also note that $\mu$ is the *expected* value of fecundity and thus *observed* values (i.e., draws from the prior predictive distribution) can be much larger (or smaller), especially as $\phi$ approaches 0.

We also considered a model in which (1) a negative binomial likelihood (without zero-inflation) was employed. Additionally, we considered models in which Equation 13 was simplified as follows: (2) The expected probability of zero-inflation ($\theta$) varies by species, including $\gamma_\theta$ that varies by species and site. (3) $\theta$ varies by species and treatment, including $\gamma_\theta$ that varies by species, treatment, and site. We used LOO-CV to compare alternative models and found that all zero-inflated models were estimated to have better predictive performance than model (1) (Table S3.12), reflecting an excess of zeros in our fecundity data. All zero-inflated models were estimated to have similar predictive performance, especially model (3) and the maximal model (described above). This is unsurprising, as the former represents variation in $\theta$ across sites entirely with $\gamma_\theta$, whereas the latter instead represents some of this variation with $\beta_{\theta,j}$. Therefore, we used the maximal model for our analyses, as it better aligns with our objective of quantifying demographic responses to environmental gradients. We followed the same procedure for fitting, checking, and comparing these models as described for the germination model.

**Fitness model**

We observed bimodalities in empirical estimates of fitness computed from paired measurements of germination rate and fecundity in each subplot, likely due to the excess of zeros in our fecundity data. Therefore, rather than modeling fitness directly, we implemented a joint model of the germination and fecundity models described above. This joint modeling approach allowed us to compute fitness ($r$) using the joint posterior distribution for the germination and fecundity sub-models. We followed the same procedure for fitting and checking this model as described for the individual models.

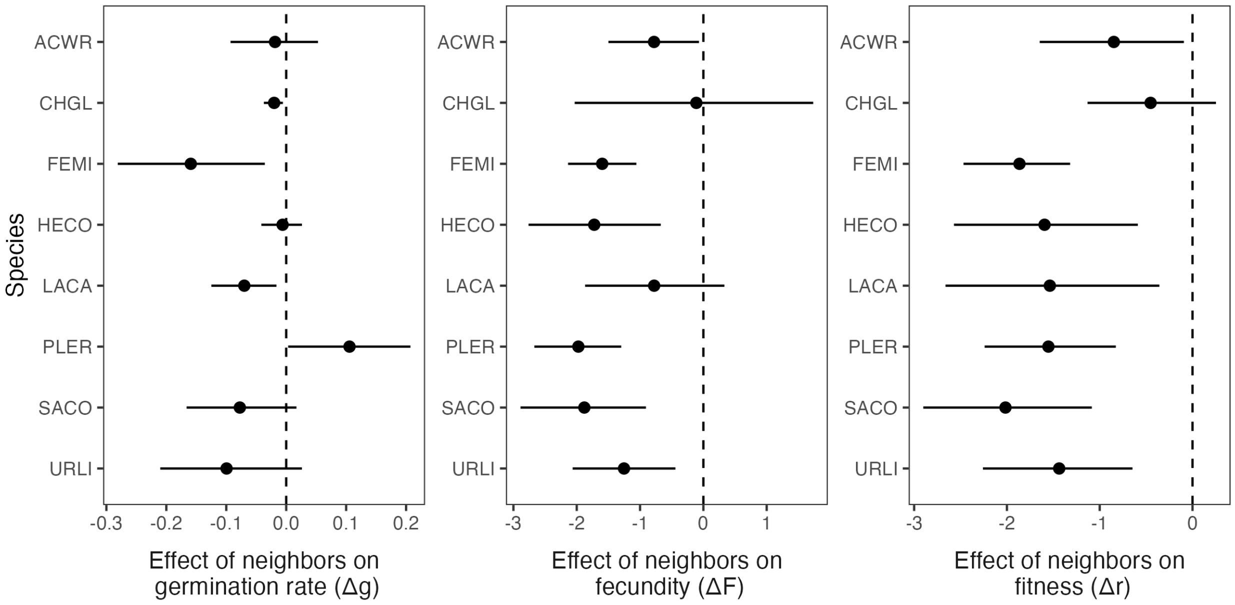

Figure S3.1: Estimated effects of neighbors on germination rate ($\Delta g$), fecundity ($\Delta F$), and fitness ($\Delta r$). For each demographic quantity ($y$), $\Delta y = y_{\text{trt}=1} - y_{\text{trt}=2}$ where trt $= 1$ is with neighbors present and trt $= 2$ is with neighbors absent. Thus, negative values of $\Delta y$ correspond to negative effects of neighbors. For each species, effects are computed on the logit scale for germination rate, log scale for fecundity, and identity scale for fitness, with soil PC1–PC4 held at average conditions across experimental sites. Points represent posterior medians. Lines represent 95% highest-density continuous intervals.

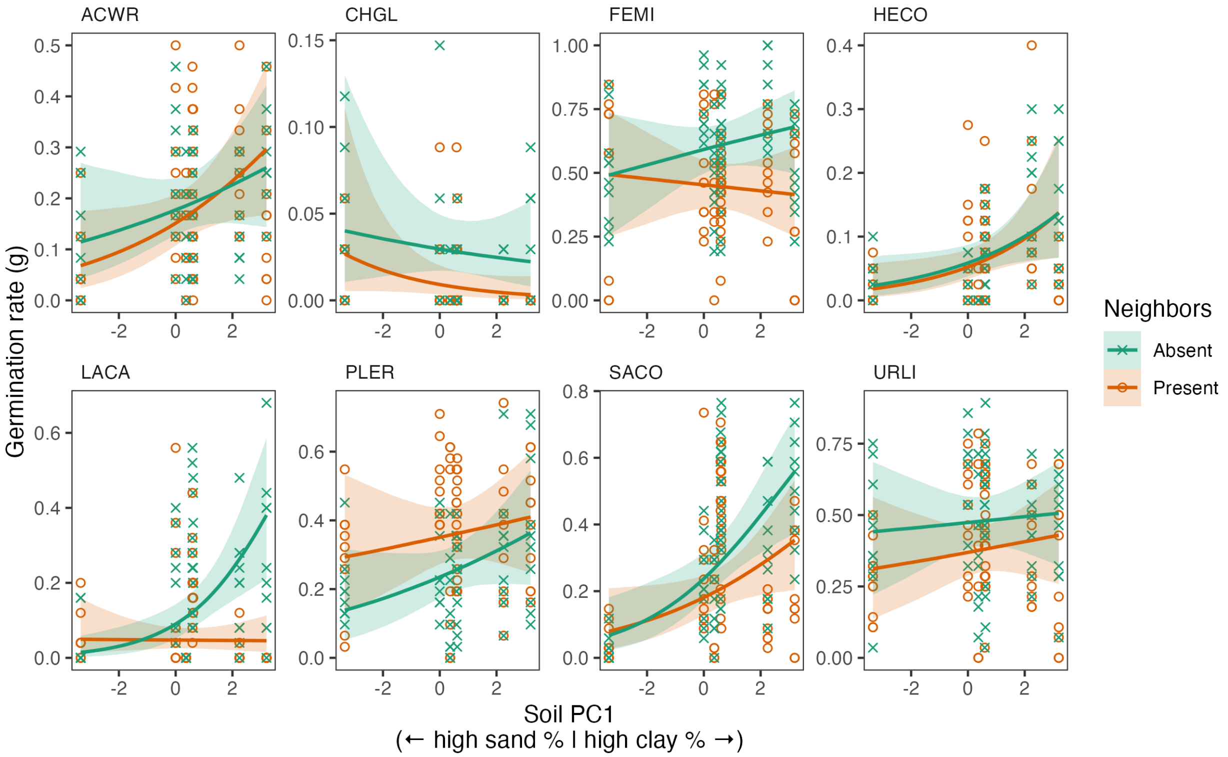

Figure S3.2: Response of germination rate ($g$) to soil PC1 in the presence or absence of neighbors. Posterior expectations are shown for each species with soil PC2–PC4 held at average conditions across experimental sites. Solid lines represent medians. Shaded areas represent 95% quantile intervals.

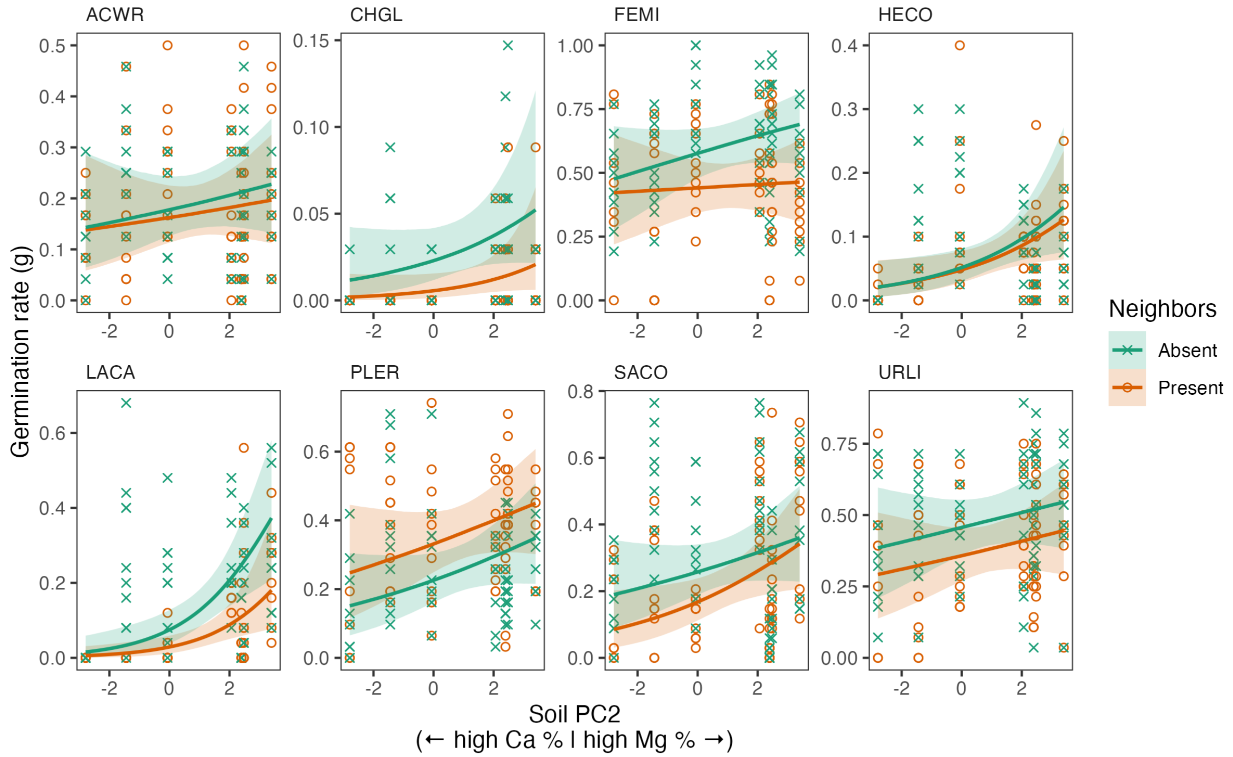

Figure S3.3: Response of germination rate ($g$) to soil PC2 in the presence or absence of neighbors. Posterior expectations are shown for each species with soil PC1, PC3, and PC4 held at average conditions across experimental sites. Solid lines represent medians. Shaded areas represent 95% quantile intervals.

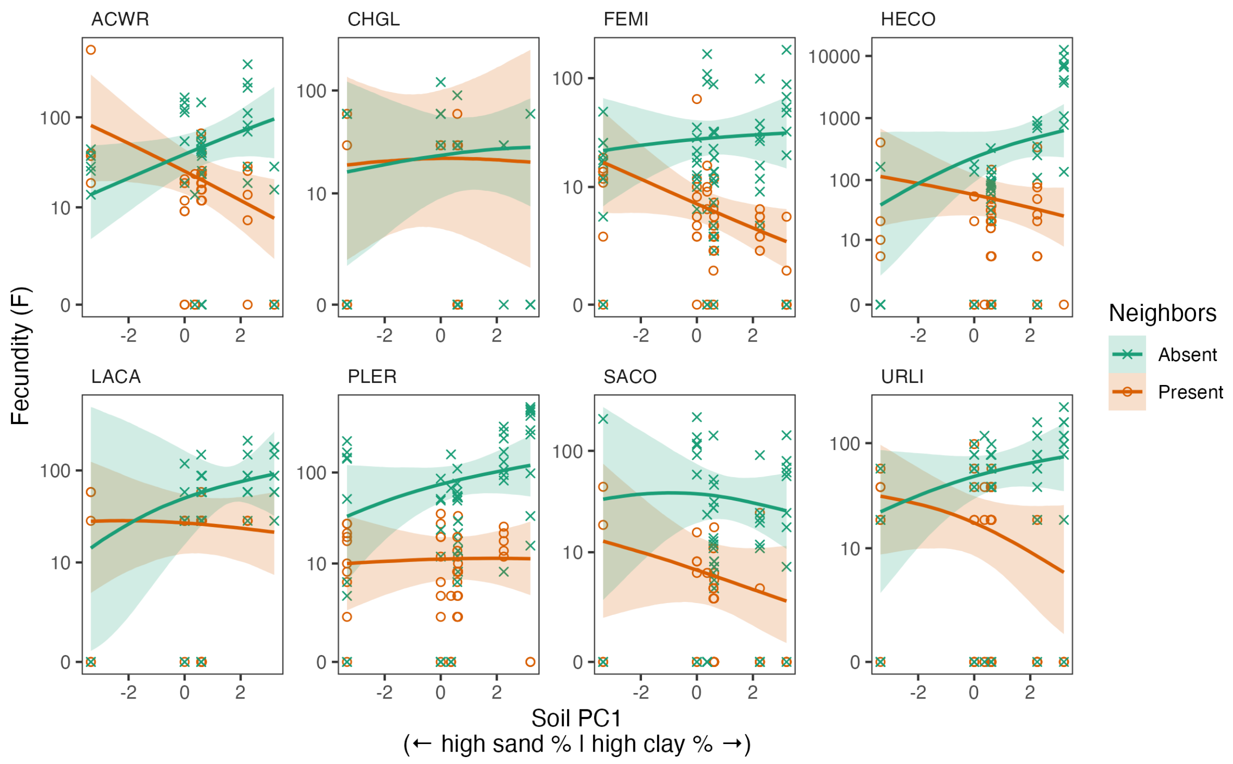

Figure S3.4: Response of fecundity ($F$) to soil PC1 in the presence or absence of neighbors. Posterior expectations are shown for each species with soil PC2–PC4 held at average conditions across experimental sites. Solid lines represent medians. Shaded areas represent 95% quantile intervals.

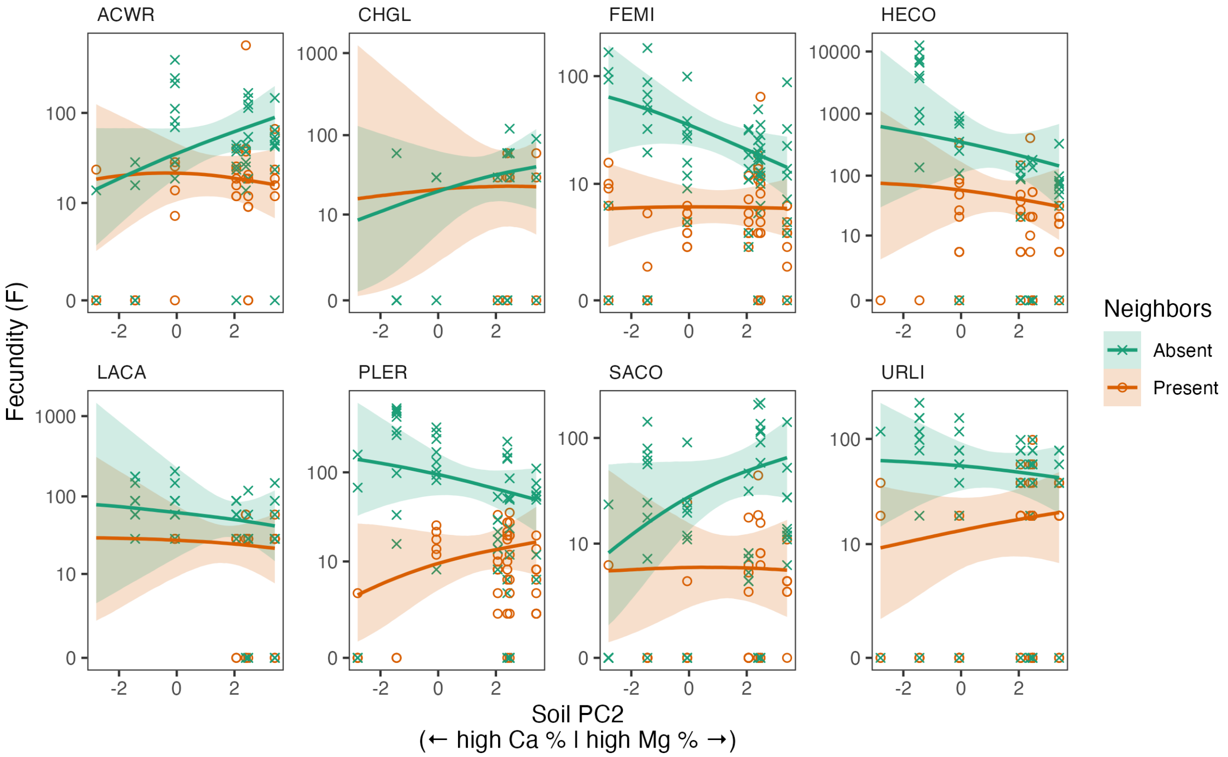

Figure S3.5: Response of fecundity ($F$) to soil PC2 in the presence or absence of neighbors. Posterior expectations are shown for each species with soil PC1, PC3, and PC4 held at average conditions across experimental sites. Solid lines represent medians. Shaded areas represent 95% quantile intervals.

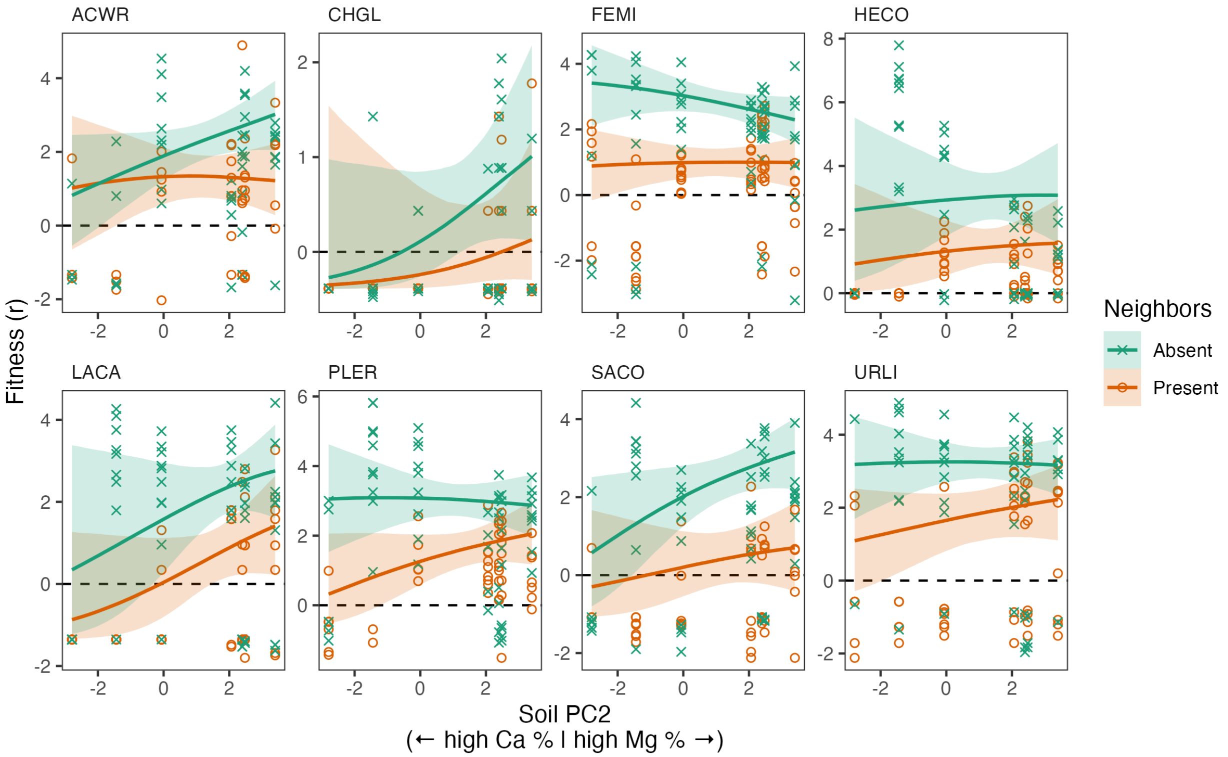

Figure S3.6: Response of fitness ($r$) to soil PC2 in the presence or absence of neighbors. Posterior expectations are shown for each species with soil PC1, PC3, and PC4 held at average conditions across experimental sites. Solid lines represent medians. Shaded areas represent 95% quantile intervals.

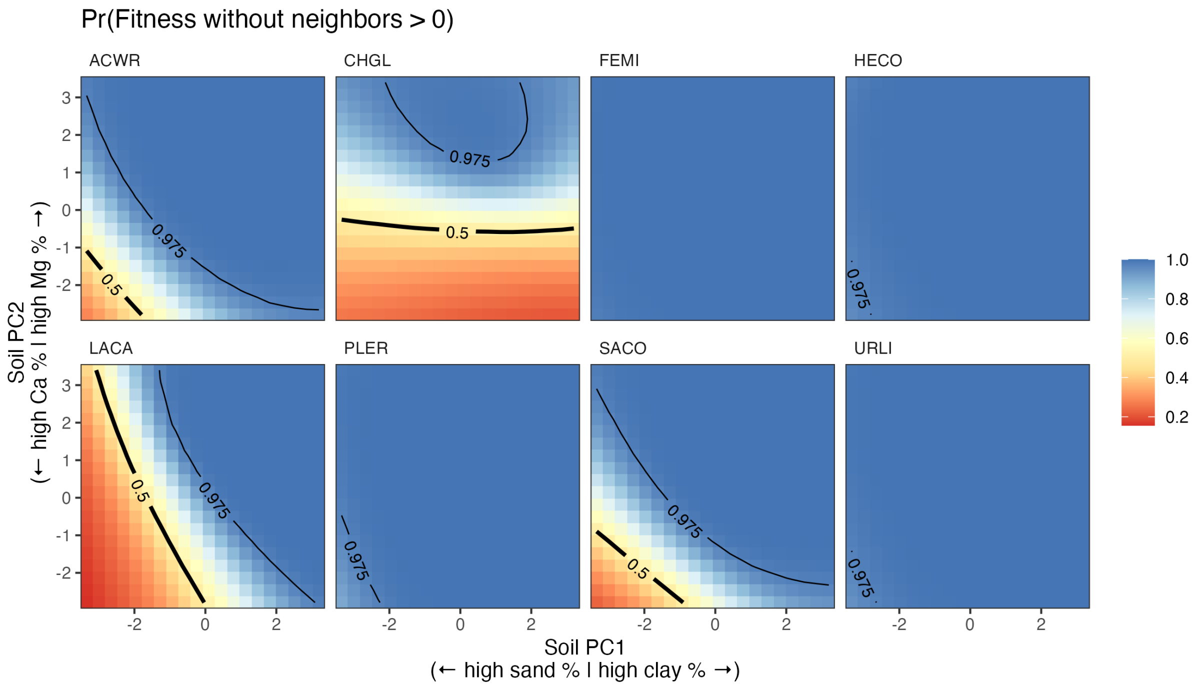

Figure S3.7: Probability that fitness ($r$) > 0 in the absence of neighbors with respect to soil PC1 and PC2. Posterior probabilities are shown for each species with soil PC3 and PC4 held at average conditions across experimental sites.

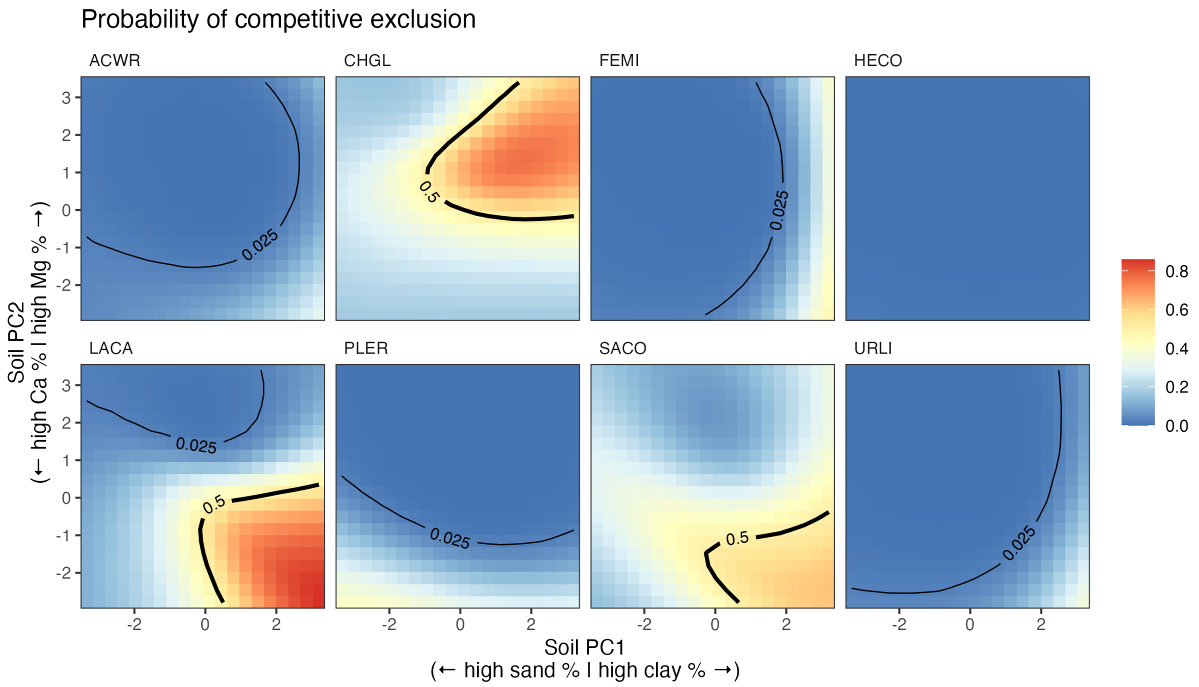

Figure S3.8: Probability of competitive exclusion (i.e., fitness ($r$) is reduced from positive in the absence of neighbors to negative in the presence of neighbors) with respect to soil PC1 and PC2. Posterior probabilities are shown for each species with soil PC3 and PC4 held at average conditions across experimental sites.

Table S3.1: Estimated effects of neighbors on germination rate ($\Delta g$), fecundity ($\Delta F$), and fitness ($\Delta r$). For each demographic quantity ($y$), $\Delta y = y_{\text{trt}=1} - y_{\text{trt}=2}$ where trt $= 1$ is with neighbors present and trt $= 2$ is with neighbors absent. Thus, negative values of $\Delta y$ correspond to negative effects of neighbors. For each species, effects are computed on the logit scale for germination rate, log scale for fecundity, and identity scale for fitness, with soil PC1–PC4 held at average conditions across experimental sites. Bold text indicates Pr(direction) $\geq$ 0.975. Italic text indicates 0.95 $\leq$ Pr(direction) < 0.975.

| **Response** | **Species** | **Median** | **95% HDCI** | **Pr(direction)** |
|---|---|---|---|---|
| Germination rate | ACWR | -0.019 | [-0.093, 0.053] | 0.702 |
| | **CHGL** | **-0.020** | **[-0.037, -0.006]** | **0.998** |
| | **FEMI** | **-0.159** | **[-0.281, -0.036]** | **0.993** |
| | HECO | -0.006 | [-0.041, 0.026] | 0.640 |
| | **LACA** | **-0.070** | **[-0.125, -0.016]** | **0.997** |
| | **PLER** | **0.106** | **[ 0.003, 0.207]** | **0.978** |
| | *SACO* | *-0.077* | *[-0.166, 0.017]* | *0.951* |
| | URLI | -0.100 | [-0.210, 0.026] | 0.949 |
| Fecundity | **ACWR** | **-0.779** | **[-1.499, -0.069]** | **0.982** |
| | CHGL | -0.111 | [-2.033, 1.735] | 0.546 |
| | **FEMI** | **-1.598** | **[-2.137, -1.058]** | **1.000** |
| | **HECO** | **-1.724** | **[-2.761, -0.673]** | **0.999** |
| | LACA | -0.777 | [-1.868, 0.332] | 0.920 |
| | **PLER** | **-1.974** | **[-2.670, -1.298]** | **1.000** |
| | **SACO** | **-1.880** | **[-2.888, -0.907]** | **1.000** |
| | **URLI** | **-1.253** | **[-2.065, -0.441]** | **0.999** |
| Fitness | **ACWR** | **-0.848** | **[-1.647, -0.093]** | **0.982** |
| | CHGL | -0.451 | [-1.131, 0.253] | 0.910 |
| | **FEMI** | **-1.865** | **[-2.468, -1.319]** | **1.000** |
| | **HECO** | **-1.595** | **[-2.571, -0.589]** | **0.999** |
| | **LACA** | **-1.537** | **[-2.662, -0.357]** | **0.995** |
| | **PLER** | **-1.553** | **[-2.240, -0.826]** | **1.000** |
| | **SACO** | **-2.016** | **[-2.901, -1.085]** | **1.000** |
| | **URLI** | **-1.437** | **[-2.258, -0.647]** | **1.000** |

Table S3.2: Estimated effects of soil PC1–PC4 on germination rate ($g$) with neighbors *absent* ($\beta_{\text{trt}=2}$). Effects are computed on the logit scale. Bold text indicates Pr(direction) $\geq$ 0.975. Italic text indicates 0.95 $\leq$ Pr(direction) < 0.975.

| Variable | Species | Median | 95% HDCI | Pr(direction) |
|---|---|---|---|---|
| PC1 | ACWR | 0.314 | [-0.185, 0.840] | 0.885 |
| | CHGL | -0.189 | [-0.871, 0.426] | 0.717 |
| | FEMI | 0.248 | [-0.275, 0.765] | 0.827 |
| | **HECO** | **0.598** | **[ 0.044, 1.164]** | **0.982** |
| | **LACA** | **1.172** | **[ 0.492, 1.871]** | **1.000** |
| | PLER | 0.400 | [-0.142, 0.894] | 0.935 |
| | **SACO** | **0.900** | **[ 0.369, 1.457]** | **0.999** |
| | URLI | 0.081 | [-0.414, 0.605] | 0.626 |
| PC2 | ACWR | 0.211 | [-0.282, 0.741] | 0.799 |
| | CHGL | 0.574 | [-0.184, 1.376] | 0.936 |
| | FEMI | 0.337 | [-0.148, 0.858] | 0.907 |
| | **HECO** | **0.782** | **[ 0.128, 1.485]** | **0.992** |
| | **LACA** | **1.360** | **[ 0.574, 2.186]** | **1.000** |
| | PLER | 0.410 | [-0.113, 0.945] | 0.940 |
| | SACO | 0.330 | [-0.161, 0.827] | 0.909 |
| | URLI | 0.244 | [-0.260, 0.746] | 0.836 |
| PC3 | ACWR | -0.056 | [-0.485, 0.366] | 0.606 |
| | CHGL | -0.059 | [-0.603, 0.478] | 0.585 |
| | FEMI | -0.102 | [-0.540, 0.333] | 0.679 |
| | HECO | -0.057 | [-0.513, 0.378] | 0.603 |
| | LACA | 0.200 | [-0.241, 0.650] | 0.815 |
| | PLER | -0.216 | [-0.657, 0.203] | 0.841 |
| | **SACO** | **0.490** | **[ 0.079, 0.896]** | **0.990** |
| | URLI | -0.027 | [-0.443, 0.398] | 0.553 |
| PC4 | ACWR | -0.123 | [-0.698, 0.443] | 0.673 |
| | CHGL | -0.356 | [-1.104, 0.382] | 0.832 |
| | FEMI | -0.154 | [-0.714, 0.469] | 0.697 |
| | HECO | -0.438 | [-1.055, 0.181] | 0.922 |
| | LACA | -0.399 | [-1.064, 0.245] | 0.888 |
| | PLER | -0.138 | [-0.714, 0.440] | 0.684 |
| | SACO | -0.047 | [-0.609, 0.497] | 0.569 |
| | URLI | -0.067 | [-0.626, 0.521] | 0.592 |

Table S3.3: Estimated effects of soil PC1–PC4 on germination rate ($g$) with neighbors *present* ($\beta_{\mathrm{trt}=1}$). Effects are computed on the logit scale. Bold text indicates Pr(direction) $\geq$ 0.975. Italic text indicates 0.95 $\leq$ Pr(direction) < 0.975.

| Variable | Species | Median | 95% HDCI | Pr(direction) |
|---|---|---|---|---|
| PC1 | **ACWR** | **0.553** | **[ 0.033, 1.082]** | **0.980** |
| | CHGL | -0.677 | [-1.536, 0.137] | 0.947 |
| | FEMI | -0.101 | [-0.628, 0.425] | 0.650 |
| | **HECO** | **0.681** | **[ 0.100, 1.257]** | **0.989** |
| | LACA | -0.027 | [-0.676, 0.608] | 0.535 |
| | PLER | 0.162 | [-0.337, 0.685] | 0.740 |
| | **SACO** | **0.580** | **[ 0.047, 1.125]** | **0.981** |
| | URLI | 0.160 | [-0.358, 0.676] | 0.731 |
| PC2 | ACWR | 0.158 | [-0.404, 0.672] | 0.728 |
| | CHGL | 0.900 | [-0.205, 2.160] | 0.948 |
| | FEMI | 0.062 | [-0.472, 0.595] | 0.591 |
| | **HECO** | **0.714** | **[ 0.063, 1.390]** | **0.987** |
| | **LACA** | **1.356** | **[ 0.443, 2.414]** | **0.999** |
| | PLER | 0.342 | [-0.160, 0.861] | 0.908 |
| | **SACO** | **0.645** | **[ 0.067, 1.234]** | **0.986** |
| | URLI | 0.253 | [-0.262, 0.806] | 0.826 |
| PC3 | ACWR | -0.120 | [-0.547, 0.309] | 0.717 |
| | CHGL | 0.192 | [-0.526, 0.872] | 0.709 |
| | FEMI | 0.060 | [-0.378, 0.486] | 0.610 |
| | HECO | -0.093 | [-0.545, 0.347] | 0.664 |
| | *LACA* | *0.420* | *[-0.058, 0.908]* | *0.955* |
| | PLER | -0.016 | [-0.419, 0.395] | 0.530 |
| | **SACO** | **0.440** | **[ 0.030, 0.859]** | **0.981** |
| | URLI | 0.067 | [-0.356, 0.489] | 0.620 |
| PC4 | ACWR | 0.184 | [-0.386, 0.765] | 0.738 |
| | CHGL | 0.019 | [-0.939, 0.966] | 0.515 |
| | FEMI | -0.041 | [-0.648, 0.534] | 0.552 |
| | HECO | 0.133 | [-0.496, 0.750] | 0.664 |
| | LACA | 0.230 | [-0.510, 0.981] | 0.728 |
| | PLER | -0.171 | [-0.730, 0.419] | 0.725 |
| | SACO | 0.085 | [-0.493, 0.675] | 0.621 |
| | URLI | 0.080 | [-0.519, 0.655] | 0.607 |

Table S3.4: Estimated effects of neighbors on the response of germination rate ($g$) to soil PC1–PC4 ($\Delta\beta = \beta_{\text{trt}=1} - \beta_{\text{trt}=2}$). Effects are computed on the logit scale. Bold text indicates Pr(direction) $\geq 0.975$. Italic text indicates $0.95 \leq$ Pr(direction) < 0.975.

| Variable | Species | Median | 95% HDCI | Pr(direction) |
|---|---|---|---|---|
| PC1 | ACWR | 0.239 | [-0.497, 0.954] | 0.740 |
| | CHGL | -0.490 | [-1.528, 0.585] | 0.821 |
| | FEMI | -0.344 | [-1.078, 0.398] | 0.825 |
| | HECO | 0.088 | [-0.715, 0.884] | 0.584 |
| | **LACA** | **-1.207** | **[-2.168, -0.290]** | **0.994** |
| | PLER | -0.236 | [-0.949, 0.499] | 0.743 |
| | SACO | -0.321 | [-1.108, 0.427] | 0.802 |
| | URLI | 0.077 | [-0.698, 0.774] | 0.583 |
| PC2 | ACWR | -0.050 | [-0.792, 0.687] | 0.557 |
| | CHGL | 0.330 | [-1.094, 1.756] | 0.678 |
| | FEMI | -0.274 | [-0.995, 0.465] | 0.771 |
| | HECO | -0.064 | [-1.059, 0.874] | 0.553 |
| | LACA | 0.004 | [-1.229, 1.309] | 0.502 |
| | PLER | -0.071 | [-0.802, 0.672] | 0.575 |
| | SACO | 0.317 | [-0.463, 1.067] | 0.794 |
| | URLI | 0.005 | [-0.700, 0.776] | 0.506 |
| PC3 | ACWR | -0.065 | [-0.672, 0.544] | 0.586 |
| | CHGL | 0.248 | [-0.634, 1.118] | 0.714 |
| | FEMI | 0.163 | [-0.474, 0.760] | 0.700 |
| | HECO | -0.038 | [-0.674, 0.589] | 0.550 |
| | LACA | 0.221 | [-0.423, 0.886] | 0.746 |
| | PLER | 0.201 | [-0.407, 0.795] | 0.751 |
| | SACO | -0.051 | [-0.623, 0.533] | 0.572 |
| | URLI | 0.092 | [-0.505, 0.694] | 0.623 |
| PC4 | ACWR | 0.307 | [-0.500, 1.123] | 0.779 |
| | CHGL | 0.375 | [-0.788, 1.630] | 0.728 |
| | FEMI | 0.114 | [-0.723, 0.937] | 0.606 |
| | HECO | 0.573 | [-0.286, 1.470] | 0.902 |
| | LACA | 0.624 | [-0.353, 1.640] | 0.896 |
| | PLER | -0.029 | [-0.876, 0.751] | 0.529 |
| | SACO | 0.139 | [-0.687, 0.919] | 0.634 |
| | URLI | 0.149 | [-0.680, 0.979] | 0.639 |

Table S3.5: Estimated effects of soil PC1–PC4 on fecundity ($F$) with neighbors *absent* ($\beta^*_{\text{trt}=2}$). Effects are computed on the log scale. Bold text indicates Pr(direction) $\geq$ 0.975. Italic text indicates 0.95 $\leq$ Pr(direction) < 0.975.

| Variable | Species | Median | 95% HDCI | Pr(direction) |
|---|---|---|---|---|
| PC1 | *ACWR* | *0.604* | *[-0.022, 1.163]* | *0.972* |
| | CHGL | 0.174 | [-0.766, 1.106] | 0.650 |
| | FEMI | 0.119 | [-0.414, 0.669] | 0.677 |
| | HECO | 0.884 | [-0.179, 2.002] | 0.949 |
| | LACA | 0.583 | [-0.809, 1.997] | 0.801 |
| | PLER | 0.400 | [-0.201, 1.040] | 0.918 |
| | SACO | -0.076 | [-0.934, 0.836] | 0.570 |
| | URLI | 0.382 | [-0.242, 1.056] | 0.894 |
| PC2 | *ACWR* | *0.689* | *[-0.129, 1.484]* | *0.950* |
| | CHGL | 0.599 | [-0.802, 2.102] | 0.800 |
| | *FEMI* | *-0.560* | *[-1.129, 0.056]* | *0.966* |
| | HECO | -0.548 | [-2.131, 1.004] | 0.753 |
| | LACA | -0.234 | [-1.655, 1.197] | 0.628 |
| | PLER | -0.377 | [-1.166, 0.350] | 0.840 |
| | SACO | 0.794 | [-0.116, 1.801] | 0.948 |
| | URLI | -0.136 | [-0.830, 0.545] | 0.660 |
| PC3 | **ACWR** | **-0.528** | **[-1.036, -0.025]** | **0.981** |
| | CHGL | -0.316 | [-1.249, 0.566] | 0.771 |
| | FEMI | -0.068 | [-0.521, 0.379] | 0.628 |
| | HECO | -0.264 | [-0.943, 0.452] | 0.770 |
| | LACA | 0.120 | [-0.351, 0.612] | 0.708 |
| | **PLER** | **-0.512** | **[-0.956, -0.049]** | **0.983** |
| | SACO | -0.398 | [-0.939, 0.177] | 0.913 |
| | URLI | 0.087 | [-0.351, 0.532] | 0.666 |
| PC4 | ACWR | -0.042 | [-0.731, 0.633] | 0.548 |
| | CHGL | -0.040 | [-1.049, 0.986] | 0.532 |
| | FEMI | 0.157 | [-0.444, 0.743] | 0.704 |
| | HECO | -0.590 | [-1.608, 0.410] | 0.876 |
| | LACA | -0.029 | [-0.780, 0.715] | 0.532 |
| | PLER | 0.014 | [-0.657, 0.645] | 0.517 |
| | SACO | -0.453 | [-1.178, 0.297] | 0.888 |
| | URLI | -0.059 | [-0.666, 0.579] | 0.580 |

Table S3.6: Estimated effects of soil PC1–PC4 on fecundity ($F$) with neighbors *present* ($\beta^{*}_{\text{trt}=1}$). Effects are computed on the log scale. Bold text indicates Pr(direction) $\geq$ 0.975. Italic text indicates 0.95 $\leq$ Pr(direction) < 0.975.

| Variable | Species | Median | 95% HDCI | Pr(direction) |
|---|---|---|---|---|
| PC1 | **ACWR** | **-0.760** | **[-1.429, -0.090]** | **0.985** |
| | CHGL | 0.026 | [-1.057, 1.199] | 0.518 |
| | **FEMI** | **-0.593** | **[-1.142, -0.035]** | **0.981** |
| | HECO | -0.472 | [-1.257, 0.348] | 0.883 |
| | LACA | -0.080 | [-0.755, 0.639] | 0.596 |
| | PLER | 0.038 | [-0.565, 0.658] | 0.550 |
| | SACO | -0.490 | [-1.406, 0.446] | 0.853 |
| | URLI | -0.550 | [-1.370, 0.250] | 0.914 |
| PC2 | ACWR | -0.042 | [-1.014, 0.894] | 0.534 |
| | CHGL | 0.130 | [-1.777, 2.104] | 0.555 |
| | FEMI | 0.003 | [-0.585, 0.557] | 0.505 |
| | HECO | -0.329 | [-1.761, 1.216] | 0.666 |
| | LACA | -0.116 | [-1.319, 1.174] | 0.571 |
| | PLER | 0.564 | [-0.439, 1.514] | 0.874 |
| | SACO | 0.012 | [-1.206, 1.218] | 0.508 |
| | URLI | 0.304 | [-0.504, 1.193] | 0.773 |
| PC3 | ACWR | -0.059 | [-0.588, 0.442] | 0.593 |
| | CHGL | -0.108 | [-1.014, 0.870] | 0.596 |
| | FEMI | -0.036 | [-0.465, 0.430] | 0.564 |
| | HECO | -0.102 | [-0.688, 0.474] | 0.641 |
| | LACA | -0.051 | [-0.533, 0.442] | 0.590 |
| | PLER | -0.159 | [-0.642, 0.314] | 0.757 |
| | SACO | -0.202 | [-0.916, 0.495] | 0.726 |
| | URLI | 0.188 | [-0.472, 0.846] | 0.720 |
| PC4 | *ACWR* | *0.759* | *[-0.119, 1.722]* | *0.957* |
| | CHGL | 0.190 | [-0.958, 1.249] | 0.640 |
| | FEMI | -0.154 | [-0.843, 0.539] | 0.670 |
| | HECO | -0.155 | [-1.251, 0.928] | 0.615 |
| | LACA | 0.124 | [-0.666, 0.948] | 0.626 |
| | PLER | -0.248 | [-1.085, 0.589] | 0.725 |
| | SACO | 0.211 | [-0.857, 1.316] | 0.651 |
| | URLI | 0.056 | [-0.866, 0.997] | 0.547 |

Table S3.7: Estimated effects of neighbors on the response of fecundity ($F$) to soil PC1–PC4 ($\Delta\beta^* = \beta^*_{\text{trt}=1} - \beta^*_{\text{trt}=2}$). Effects are computed on the log scale. Bold text indicates Pr(direction) $\geq$ 0.975. Italic text indicates 0.95 $\leq$ Pr(direction) < 0.975.

| Variable | Species | Median | 95% HDCI | Pr(direction) |
|---|---|---|---|---|
| PC1 | **ACWR** | **-1.362** | **[-2.262, -0.465]** | **0.997** |
| | CHGL | -0.157 | [-1.709, 1.234] | 0.584 |
| | *FEMI* | *-0.712* | *[-1.494, 0.065]* | *0.963* |
| | **HECO** | **-1.360** | **[-2.725, -0.035]** | **0.976** |
| | LACA | -0.667 | [-2.245, 0.903] | 0.805 |
| | PLER | -0.371 | [-1.238, 0.497] | 0.806 |
| | SACO | -0.419 | [-1.649, 0.899] | 0.742 |
| | *URLI* | *-0.951* | *[-2.009, 0.091]* | *0.964* |
| PC2 | ACWR | -0.730 | [-1.980, 0.523] | 0.873 |
| | CHGL | -0.485 | [-2.889, 1.902] | 0.652 |
| | FEMI | 0.562 | [-0.261, 1.386] | 0.911 |
| | HECO | 0.231 | [-2.073, 2.270] | 0.583 |
| | LACA | 0.125 | [-1.770, 2.032] | 0.554 |
| | PLER | 0.945 | [-0.259, 2.204] | 0.933 |
| | SACO | -0.789 | [-2.274, 0.789] | 0.841 |
| | URLI | 0.449 | [-0.615, 1.530] | 0.798 |
| PC3 | ACWR | 0.471 | [-0.254, 1.198] | 0.902 |
| | CHGL | 0.219 | [-1.073, 1.526] | 0.635 |
| | FEMI | 0.030 | [-0.597, 0.668] | 0.540 |
| | HECO | 0.158 | [-0.768, 1.043] | 0.635 |
| | LACA | -0.173 | [-0.848, 0.524] | 0.713 |
| | PLER | 0.353 | [-0.322, 1.010] | 0.852 |
| | SACO | 0.183 | [-0.707, 1.076] | 0.657 |
| | URLI | 0.096 | [-0.736, 0.853] | 0.602 |
| PC4 | ACWR | 0.795 | [-0.334, 1.953] | 0.917 |
| | CHGL | 0.239 | [-1.217, 1.725] | 0.625 |
| | FEMI | -0.313 | [-1.208, 0.626] | 0.751 |
| | HECO | 0.440 | [-1.017, 1.939] | 0.723 |
| | LACA | 0.148 | [-0.929, 1.250] | 0.615 |
| | PLER | -0.256 | [-1.291, 0.824] | 0.690 |
| | SACO | 0.664 | [-0.635, 1.997] | 0.842 |
| | URLI | 0.113 | [-0.974, 1.274] | 0.583 |

Table S3.8: Estimated effects of soil PC1–PC4 on fitness ($r$) with neighbors *absent* ($\beta^*_{\text{trt}=2}$). Effects are computed on the identity scale. Bold text indicates Pr(direction) $\geq$ 0.975. Italic text indicates 0.95 $\leq$ Pr(direction) < 0.975.

| Variable | Species | Median | 95% HDCI | Pr(direction) |
|---|---|---|---|---|
| PC1 | **ACWR** | **0.817** | **[ 0.143, 1.430]** | **0.987** |
| | CHGL | -0.001 | [-0.598, 0.465] | 0.501 |
| | FEMI | 0.223 | [-0.344, 0.816] | 0.784 |
| | **HECO** | **1.194** | **[ 0.347, 1.768]** | **0.991** |
| | **LACA** | **1.346** | **[ 0.292, 1.960]** | **0.984** |
| | **PLER** | **0.675** | **[ 0.031, 1.311]** | **0.977** |
| | SACO | 0.550 | [-0.266, 1.308] | 0.889 |
| | URLI | 0.422 | [-0.230, 1.103] | 0.901 |
| PC2 | *ACWR* | *0.828* | *[ 0.000, 1.625]* | *0.968* |
| | CHGL | 0.461 | [-0.178, 0.986] | 0.926 |
| | FEMI | -0.416 | [-1.034, 0.219] | 0.905 |
| | HECO | 0.172 | [-1.289, 1.596] | 0.586 |
| | LACA | 0.894 | [-0.432, 1.936] | 0.887 |
| | PLER | -0.065 | [-0.899, 0.765] | 0.563 |
| | **SACO** | **0.976** | **[ 0.101, 1.776]** | **0.975** |
| | URLI | -0.004 | [-0.714, 0.734] | 0.504 |
| PC3 | *ACWR* | *-0.554* | *[-1.114, 0.008]* | *0.972* |
| | CHGL | -0.167 | [-0.557, 0.342] | 0.773 |
| | FEMI | -0.109 | [-0.596, 0.369] | 0.684 |
| | HECO | -0.298 | [-1.002, 0.461] | 0.781 |
| | LACA | 0.283 | [-0.299, 0.866] | 0.838 |
| | **PLER** | **-0.650** | **[-1.149, -0.110]** | **0.991** |
| | SACO | -0.068 | [-0.643, 0.536] | 0.587 |
| | URLI | 0.069 | [-0.416, 0.548] | 0.619 |
| PC4 | ACWR | -0.134 | [-0.898, 0.669] | 0.631 |
| | CHGL | -0.177 | [-0.789, 0.409] | 0.725 |
| | FEMI | 0.095 | [-0.552, 0.709] | 0.620 |
| | *HECO* | *-0.922* | *[-1.824, 0.105]* | *0.956* |
| | LACA | -0.357 | [-1.176, 0.492] | 0.791 |
| | PLER | -0.086 | [-0.848, 0.648] | 0.587 |
| | SACO | -0.470 | [-1.246, 0.332] | 0.876 |
| | URLI | -0.091 | [-0.756, 0.583] | 0.610 |

Table S3.9: Estimated effects of soil PC1–PC4 on fitness ($r$) with neighbors *present* ($\beta^*_{\text{trt}=1}$). Effects are computed on the identity scale. Bold text indicates Pr(direction) $\geq$ 0.975. Italic text indicates 0.95 $\leq$ Pr(direction) < 0.975.

| Variable | Species | Median | 95% HDCI | Pr(direction) |
|---|---|---|---|---|
| PC1 | ACWR | -0.282 | [-1.008, 0.438] | 0.775 |
| | CHGL | -0.121 | [-0.669, 0.227] | 0.809 |
| | **FEMI** | **-0.610** | **[-1.184, -0.035]** | **0.978** |
| | HECO | 0.112 | [-0.590, 0.701] | 0.630 |
| | LACA | -0.086 | [-0.808, 0.598] | 0.591 |
| | PLER | 0.126 | [-0.461, 0.702] | 0.660 |
| | SACO | -0.041 | [-0.788, 0.665] | 0.544 |
| | URLI | -0.419 | [-1.146, 0.354] | 0.852 |
| PC2 | ACWR | 0.087 | [-0.874, 0.972] | 0.570 |
| | CHGL | 0.152 | [-0.432, 0.681] | 0.815 |
| | FEMI | 0.039 | [-0.569, 0.637] | 0.551 |
| | HECO | 0.240 | [-0.914, 1.097] | 0.663 |
| | LACA | 0.839 | [-0.144, 1.536] | 0.939 |
| | PLER | 0.649 | [-0.166, 1.299] | 0.930 |
| | SACO | 0.366 | [-0.563, 1.079] | 0.779 |
| | URLI | 0.425 | [-0.333, 1.177] | 0.853 |
| PC3 | ACWR | -0.147 | [-0.689, 0.447] | 0.699 |
| | CHGL | 0.009 | [-0.313, 0.374] | 0.545 |
| | FEMI | -0.006 | [-0.461, 0.475] | 0.510 |
| | HECO | -0.140 | [-0.611, 0.376] | 0.706 |
| | LACA | 0.277 | [-0.264, 0.797] | 0.846 |
| | PLER | -0.155 | [-0.620, 0.308] | 0.744 |
| | SACO | 0.081 | [-0.504, 0.654] | 0.604 |
| | URLI | 0.211 | [-0.423, 0.836] | 0.742 |
| PC4 | *ACWR* | *0.810* | *[-0.016, 1.513]* | *0.966* |
| | CHGL | 0.026 | [-0.361, 0.511] | 0.614 |
| | FEMI | -0.166 | [-0.847, 0.552] | 0.673 |
| | HECO | -0.026 | [-0.886, 0.715] | 0.524 |
| | LACA | 0.273 | [-0.546, 1.047] | 0.738 |
| | PLER | -0.306 | [-1.023, 0.483] | 0.778 |
| | SACO | 0.197 | [-0.625, 0.877] | 0.682 |
| | URLI | 0.096 | [-0.742, 0.962] | 0.583 |

Table S3.10: Estimated effects of neighbors on the response of fitness ($r$) to soil PC1–PC4 ($\Delta\beta^* = \beta^*_{\text{trt}=1} - \beta^*_{\text{trt}=2}$). Effects are computed on the identity scale. Bold text indicates Pr(direction) $\geq 0.975$. Italic text indicates $0.95 \leq$ Pr(direction) < 0.975.

| Variable | Species | Median | 95% HDCI | Pr(direction) |
|---|---|---|---|---|
| PC1 | **ACWR** | **-1.097** | **[-2.060, -0.106]** | **0.983** |
| | CHGL | -0.138 | [-0.805, 0.608] | 0.664 |
| | *FEMI* | *-0.833* | *[-1.674, -0.030]* | *0.975* |
| | *HECO* | *-1.068* | *[-2.034, -0.038]* | *0.973* |
| | **LACA** | **-1.387** | **[-2.432, -0.142]** | **0.978** |
| | PLER | -0.546 | [-1.450, 0.288] | 0.892 |
| | SACO | -0.590 | [-1.671, 0.545] | 0.847 |
| | URLI | -0.849 | [-1.845, 0.205] | 0.944 |
| PC2 | ACWR | -0.738 | [-1.985, 0.512] | 0.877 |
| | CHGL | -0.301 | [-1.076, 0.537] | 0.808 |
| | FEMI | 0.457 | [-0.421, 1.319] | 0.846 |
| | HECO | 0.036 | [-1.789, 1.816] | 0.516 |
| | LACA | -0.086 | [-1.598, 1.453] | 0.544 |
| | PLER | 0.693 | [-0.425, 1.795] | 0.883 |
| | SACO | -0.619 | [-1.861, 0.557] | 0.848 |
| | URLI | 0.430 | [-0.616, 1.467] | 0.785 |
| PC3 | ACWR | 0.408 | [-0.388, 1.216] | 0.845 |
| | CHGL | 0.175 | [-0.395, 0.714] | 0.750 |
| | FEMI | 0.103 | [-0.564, 0.772] | 0.622 |
| | HECO | 0.161 | [-0.753, 1.021] | 0.640 |
| | LACA | -0.007 | [-0.773, 0.789] | 0.506 |
| | PLER | 0.496 | [-0.220, 1.189] | 0.911 |
| | SACO | 0.144 | [-0.700, 0.963] | 0.633 |
| | URLI | 0.138 | [-0.665, 0.928] | 0.636 |
| PC4 | ACWR | 0.927 | [-0.194, 2.029] | 0.939 |
| | CHGL | 0.226 | [-0.477, 0.960] | 0.745 |
| | FEMI | -0.266 | [-1.189, 0.713] | 0.703 |
| | HECO | 0.875 | [-0.433, 2.117] | 0.903 |
| | LACA | 0.622 | [-0.554, 1.780] | 0.847 |
| | PLER | -0.216 | [-1.318, 0.816] | 0.656 |
| | SACO | 0.645 | [-0.510, 1.694] | 0.867 |
| | URLI | 0.184 | [-0.902, 1.258] | 0.630 |

Table S3.11: Comparison of germination models. We used approximate leave-one-out cross-validation (LOO-CV) to compare models with alternative likelihood functions. The model selected for use in analyses is indicated with bold text.

| **Model** | **Likelihood** | **$ELPD_{LOO}$** | **$\Delta ELPD_{LOO}$** |
|---|---|---|---|
| **(2)** | **Beta-binomial** | **-2656.74 ± 35.91** | **0.00 ± 0.00** |
| (1) | Binomial | -3361.97 ± 89.28 | -705.23 ± 67.05 |

Table S3.12: Comparison of fecundity models. We used approximate leave-one-out cross-validation (LOO-CV) to compare models with alternative likelihood functions (NB = negative binomial, ZINB = zero-inflated negative binomial) and predictors for the zero-inflation component (sp = species, trt = neighbor treatment, soil = soil PC1–PC4). The model selected for use in analyses is indicated with bold text.

| Model | Likelihood | ZI predictors | $\mathrm{ELPD}_{\mathrm{LOO}}$ | $\Delta\mathrm{ELPD}_{\mathrm{LOO}}$ |
|---|---|---|---:|---:|
| **(4)** | **ZINB** | **sp, trt, soil** | **-2849.79 ± 57.80** | **0.00 ± 0.00** |
| (3) | ZINB | sp, trt | -2851.77 ± 57.46 | -1.98 ± 3.90 |
| (2) | ZINB | sp | -2862.11 ± 57.11 | -12.32 ± 8.59 |
| (1) | NB | N/A | -3048.14 ± 53.60 | -198.35 ± 17.06 |

### Appendix S4: Analysis of occurrence surveys

We used Bayesian generalized linear models to quantify the relationship between observed occurrence (presence vs. absence) of our focal species and the soil environment. We modeled the number of plots in which presence was recorded ($n_i$) for observation $i$ = 1, …, 248 as following a beta-binomial distribution:

$$n_i \sim \text{Beta-Binomial}\left(N_i, \mu_i, \phi_i\right) \tag{21}$$

where $N$ is the number of surveyed plots per site, $\mu$ is the mean probability parameter, and $\phi$ is the precision parameter. To align the scope of this analysis with our demographic experiment, we only used occurrence data for sites within the range of soil PC1–PC4 across experimental sites, corresponding to 92 plots at 31 sites. Using a logit link function, we defined $\mu$ as:

$$\text{logit}\left(\mu_i\right) = \alpha_{\text{sp}_i} + \sum_{j=1}^{4} \beta_{j,\text{sp}_i} \times \text{PC}_{j,i} \tag{22}$$

where $\alpha$ is a species-specific intercept and $\beta_j$ is a species-specific slope for soil principal component axis $j$. Soil PC1–PC4 were centered and scaled with respect to the mean and standard deviation of each axis across sites. We allowed $\phi$ to vary by species as:

$$\phi_i = \alpha_{\phi,\text{sp}_i} \tag{23}$$

where $\alpha_\phi$ is a species-specific intercept for $\phi$.

We specified prior distributions for parameters as:

$$\alpha \sim \text{Normal}(0, 2) \tag{24}$$

$$\beta_j \sim \text{Normal}(0, 1) \tag{25}$$

These priors were specified following a similar rationale as described for the germination model in Appendix S3.

We also considered a model in which a binomial likelihood was employed. We used LOO-CV to compare alternative models and found that both models were estimated to have similar predictive performance, although $|\Delta\text{ELPD}_{\text{LOO}}|/\text{SE}_{\Delta\text{ELPD}_{\text{LOO}}}$ was close to 2 in favor of the beta-binomial model (Table S4.2). We therefore used the beta-binomial model (described above) for our analyses. We followed the same procedure for fitting, checking, and comparing these models as described for the germination model in Appendix S3.

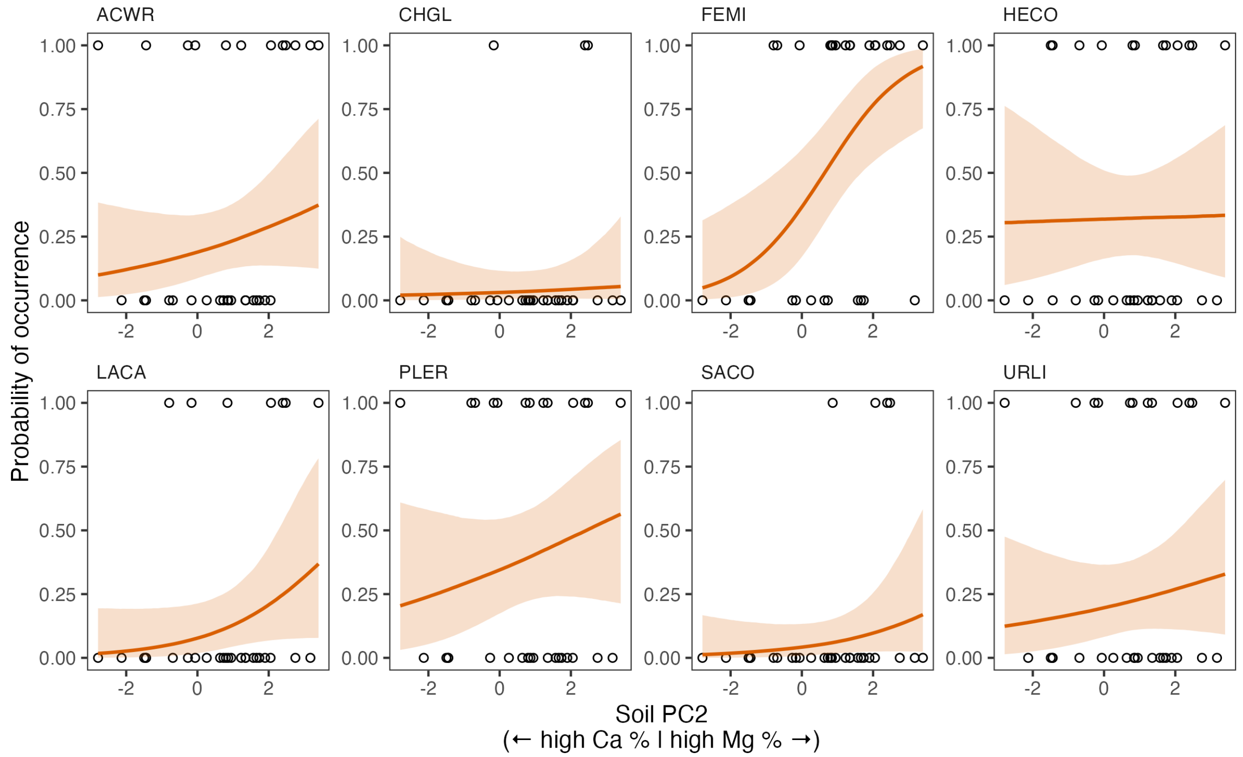

Figure S4.1: Observed patterns of occurrence along soil PC2. Posterior expectations are shown for each species with soil PC1, PC3, and PC4 held at average conditions across sites. Solid lines represent medians. Shaded areas represent 95% quantile intervals.

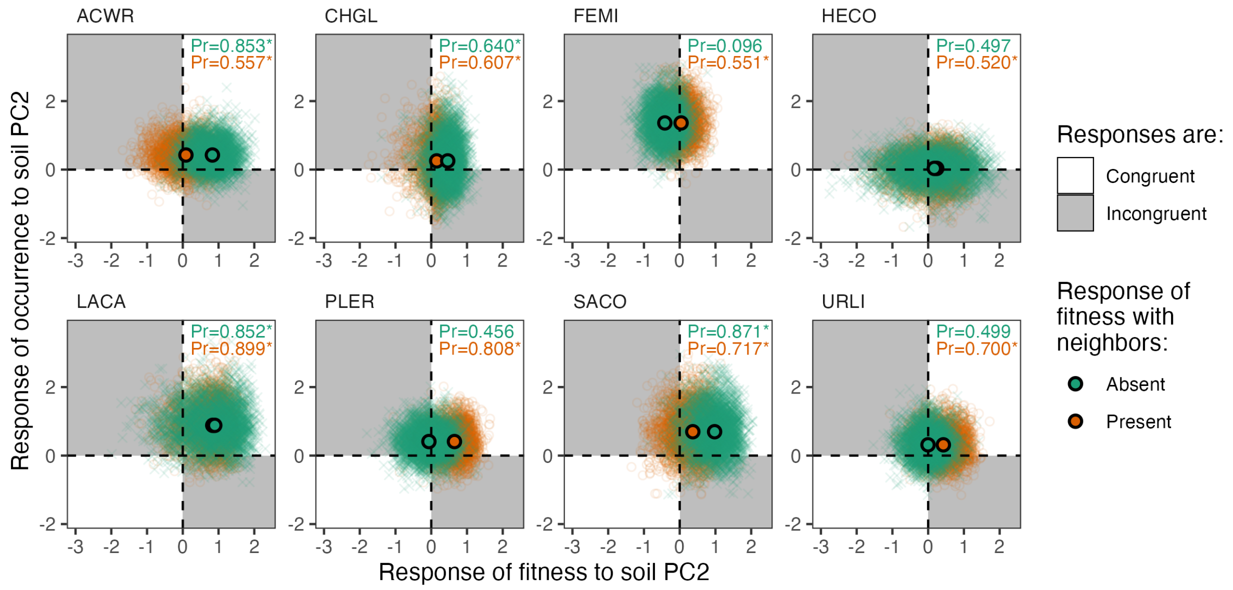

Figure S4.2: Congruence between responses of occurrence and fitness to soil PC2. For each species, the slope of occurrence in response to soil PC2 ($\beta_{j=2}$ for the occurrence model) is plotted against the axis-wide slope of fitness in response to soil PC2 ($\beta^{*}_{j=2}$ for the fitness model), the latter of which is computed in the presence (orange) or absence (green) of neighbors. Point clouds are posterior samples (thinned to 4,000 samples for visualization) and center points are medians. White regions denote congruent responses (i.e., slopes are both positive or both negative) and gray regions denote incongruent responses (i.e., one slope is positive while the other slope is negative). The total probability of congruence (i.e., the proportion of posterior samples that fall in the white regions) is shown in the top-right corner of each panel. Asterisks indicate that responses are more likely to be congruent that not (i.e., Pr(congruence) > 0.5); note that these are not results of statistical significance tests.

Table S4.1: Estimated effects of soil PC1–PC4 on occurrence. Effects are computed on the logit scale. Bold text indicates Pr(direction) $\geq$ 0.975. Italic text indicates 0.95 $\leq$ Pr(direction) < 0.975.

| Variable | Species | Median | 95% HDCI | Pr(direction) |
|---|---|---|---|---|
| PC1 | ACWR | -0.042 | [-0.689, 0.622] | 0.551 |
| | **CHGL** | **-1.241** | **[-2.273, -0.250]** | **0.993** |
| | **FEMI** | **-0.855** | **[-1.697, -0.063]** | **0.988** |
| | HECO | 0.036 | [-0.673, 0.720] | 0.541 |
| | **LACA** | **-1.076** | **[-1.977, -0.209]** | **0.994** |
| | **PLER** | **-0.873** | **[-1.684, -0.122]** | **0.991** |
| | SACO | -0.339 | [-1.225, 0.604] | 0.767 |
| | **URLI** | **-1.099** | **[-1.970, -0.275]** | **0.999** |
| PC2 | ACWR | 0.425 | [-0.298, 1.229] | 0.876 |
| | CHGL | 0.254 | [-0.885, 1.506] | 0.667 |
| | **FEMI** | **1.362** | **[ 0.449, 2.369]** | **0.999** |
| | HECO | 0.034 | [-0.772, 0.829] | 0.534 |
| | *LACA* | *0.883* | *[-0.191, 2.000]* | *0.954* |
| | PLER | 0.409 | [-0.370, 1.194] | 0.857 |
| | SACO | 0.697 | [-0.427, 1.923] | 0.890 |
| | URLI | 0.314 | [-0.470, 1.152] | 0.785 |
| PC3 | ACWR | 0.253 | [-0.373, 0.879] | 0.780 |
| | CHGL | -0.313 | [-1.387, 0.681] | 0.727 |
| | FEMI | 0.272 | [-0.515, 1.017] | 0.762 |
| | HECO | -0.015 | [-0.693, 0.629] | 0.519 |
| | **LACA** | **1.225** | **[ 0.376, 2.101]** | **0.997** |
| | **PLER** | **0.756** | **[ 0.050, 1.520]** | **0.983** |
| | SACO | 0.075 | [-0.731, 0.876] | 0.572 |
| | **URLI** | **0.767** | **[ 0.083, 1.485]** | **0.989** |
| PC4 | ACWR | -0.208 | [-0.962, 0.522] | 0.713 |
| | CHGL | -0.182 | [-1.409, 0.971] | 0.619 |
| | FEMI | -0.385 | [-1.259, 0.448] | 0.818 |
| | *HECO* | *0.762* | *[-0.009, 1.586]* | *0.974* |
| | LACA | 0.183 | [-0.727, 1.116] | 0.650 |
| | PLER | 0.038 | [-0.790, 0.815] | 0.538 |
| | SACO | -0.265 | [-1.330, 0.805] | 0.690 |
| | URLI | -0.638 | [-1.491, 0.196] | 0.939 |

Table S4.2: Comparison of occurrence models. We used approximate leave-one-out cross-validation (LOO-CV) to compare models with alternative likelihood functions. The model selected for use in analyses is indicated with bold text.

| **Model** | **Likelihood** | **$ELPD_{LOO}$** | **$\Delta ELPD_{LOO}$** |
|---|---|---|---|
| **(2)** | **Beta-binomial** | **-192.62 ± 15.73** | **0.00 ± 0.00** |
| (1) | Binomial | -210.70 ± 20.02 | -18.09 ± 10.76 |
